# Ewsr1b, Syncrip, HuR and alternative 3′UTRs organize sequential waves of translation to drive embryonic development

**DOI:** 10.1101/2025.09.18.676998

**Authors:** Keisuke Sato, Ludivine Fierro, Anyu Suginishi, Tomoya Kotani

## Abstract

Eggs of many species accumulate thousands of dormant mRNAs that are translated after fertilization at specific times and locations to direct development. However, how embryos coordinate translation of these mRNAs remains unclear. In this study, we identified sequential waves of translation critical for proper development progression. The first wave occurred within 1 h and included translation of *ewsr1b* mRNA that harbored a short 3′ untranslated region (UTR) comprising 16 nucleotides. The resulting Ewsr1b protein triggered the second translation wave through binding cytoplasmic mRNAs, including *pou5f3*, which encodes a transcription factor promoting zygotic genome activation. In contrast, HuR and Syncrip repressed translation until the first and second waves, respectively. *ewsr1b* mRNA that had a long 3′UTR was translated in the second wave, and the 3′UTR’s length determined protein localization and function. Overall, our findings reveal previously unknown molecular principles that coordinate translation timings and protein functions to drive long-term, multilayered processes.

## INTRODUCTION

Many proteins must be synthesized at the appropriate times and locations for diverse biological processes to progress. Posttranscriptional regulation, such as translational control of localized mRNAs, has emerged as a key mechanism to direct protein synthesis with temporal and spatial precision [1–3]. The best-studied examples include translation of *β-actin* mRNA at the cell protrusions during vertebrate cell migration [4–6] and translation of *ASH1* mRNA at the bud during mating-type switching in budding yeast [7–9]. In these cases, multiple proteins bind to the 3′ untranslated region (UTR) of mRNA to transport it and repress its translation until it reaches final destination. These foundational studies highlighted the importance and mechanisms of translational control in cellular processes. However, how molecular and cellular systems coordinate the timing and localization of mRNA translation in long-term, multilayered processes, such as cell differentiation, body patterning, and neuronal plasticity, remains unclear.

Embryos of many animals synthesize proteins at specific times and locations after fertilization to regulate cellular and developmental processes. In all animals studied to date, transcription is silent from shortly before fertilization until zygotic genome activation (ZGA), which occurs several to >10 h after fertilization [10–13]. Thus, temporal and spatial control of protein synthesis postfertilization relies on translational activation of dormant mRNAs stored in eggs. Previous large-scale studies of mRNA translation showed that zebrafish and mouse eggs accumulate thousands of dormant mRNAs that are translated at distinct times after fertilization, even beyond ZGA [14–17]. However, how embryos coordinate the translation of these dormant mRNAs temporally and spatially after fertilization to drive developmental processes remains largely unknown.

Zebrafish *pou5f3* mRNA encodes the transcription factor Pou5f3 (also known as Pou2/Pou5f1), which regulates multiple developmental processes after fertilization including ZGA induction, endoderm differentiation, and dorsal–ventral specification [18–22]. As a paradigm to understand temporal mRNA translation, we previously studied the regulation of *pou5f3* mRNA and found that it forms cytoplasmic RNA granules in a dormant state in oocytes, exhibiting solid-like properties [23]. Translation of this mRNA was activated after fertilization through several marked changes. First, the RNA granules became liquid droplets, enabling efficient translation within them [23]. Second, the mRNA’s 3′ end sequences were shortened before activation [24]. This shortening serves as a molecular switch by altering the proteins that bind to the 3′UTR. Although protein-binding changes were expected to alter RNA granule states, this effect remained unclear. Synaptotagmin-binding cytoplasmic RNA-interacting protein (Syncrip) and Ewing’s sarcoma RNA binding protein 1b (Ewsr1b) were identified as proteins that bind to the full-length and shortened 3′UTRs of *pou5f3* mRNA, respectively [24].

Syncrip, a member of the heterogenous nuclear ribonucleoprotein (hnRNP) family (also known as hnRNP Q), is highly conserved across organisms from fly to human [25–27]. Previous studies showed that cytoplasmic Syncrip binds the 3′UTR of mRNAs and plays key roles in posttranscriptional regulation of neuronal mRNAs [28–30]. Ewsr1b is part of the FET protein family, which includes Fus, Ewsr1, and Taf15. FET proteins share ∼70% amino acid sequence identity and are evolutionarily conserved from nematode to human [31]. A key feature of these proteins is their intrinsically disordered regions (IDRs), which contribute to liquid–liquid phase separation [32–34]. FET proteins are predominantly nuclear and regulate RNA processes, such as transcription and splicing [35–37]. Conversely, mutations in FET genes can cause cytoplasm protein aggregation, leading to neurodegenerative diseases, such as amyotrophic lateral sclerosis (ALS) [37, 38]. However, the roles of FET proteins in translational regulation, and of Syncrip and Ewsr1 in early development, remain unknown.

In this study, we found that Syncrip and Ewsr1b oppositely regulate maternal mRNAs to drive early development. Ewsr1b was synthesized immediately after fertilization from mRNA harboring a short 3′UTR. Ewsr1b then triggered changes in *pou5f3* RNA granules and translation of mRNAs, including *pou5f3*. In contrast, Syncrip was already present in eggs and repressed *pou5f3* mRNA translation. Ewsr1b was also synthesized from mRNA harboring a long 3′UTR at a time similar to *pou5f3* mRNA translation, promoting later developmental stages. The timing of Ewsr1b synthesis and its function were determined by the mRNA 3′UTR’s length. Overall, our findings reveal molecular and cellular principles for coordinating mRNA translation and protein function to drive long-term multilayered processes.

## RESULTS

### Syncrip levels increase in eggs and decline after fertilization

Our previous findings showed that dormant *pou5f3* mRNA forms cytoplasmic granules in a solid-like state in eggs, half of which transition to a liquid-like state by 3 h post fertilization (hpf) [23] (Figure 1A). During this transition, ∼70 nucleotides at the 3′ end of the mRNA are shortened, promoting translational activation by altering bound proteins [24]. Syncrip was identified as a protein that predominantly binds the full-length 3′UTR of *pou5f3* mRNA in vitro [24] (Figure 1A). Syncrip contains three sequence-specific RNA-binding domains and several IDRs (Figure 1B). To investigate Syncrip’s function, we first examined the expression patterns of *syncrip* mRNA and Syncrip protein in early zebrafish embryos. Time-course analysis revealed that *syncrip* mRNA and Syncrip protein are present in eggs, with levels gradually decreasing from 0 to 6 hpf (Figure 1C; refer to Figure S1A for mRNA). Immunofluorescence showed that Syncrip localized in cytoplasmic foci at 0 hpf (Figure 1D), and the number of foci decreased at 3 and 6 hpf (Figure S1B).

**Figure 1.**
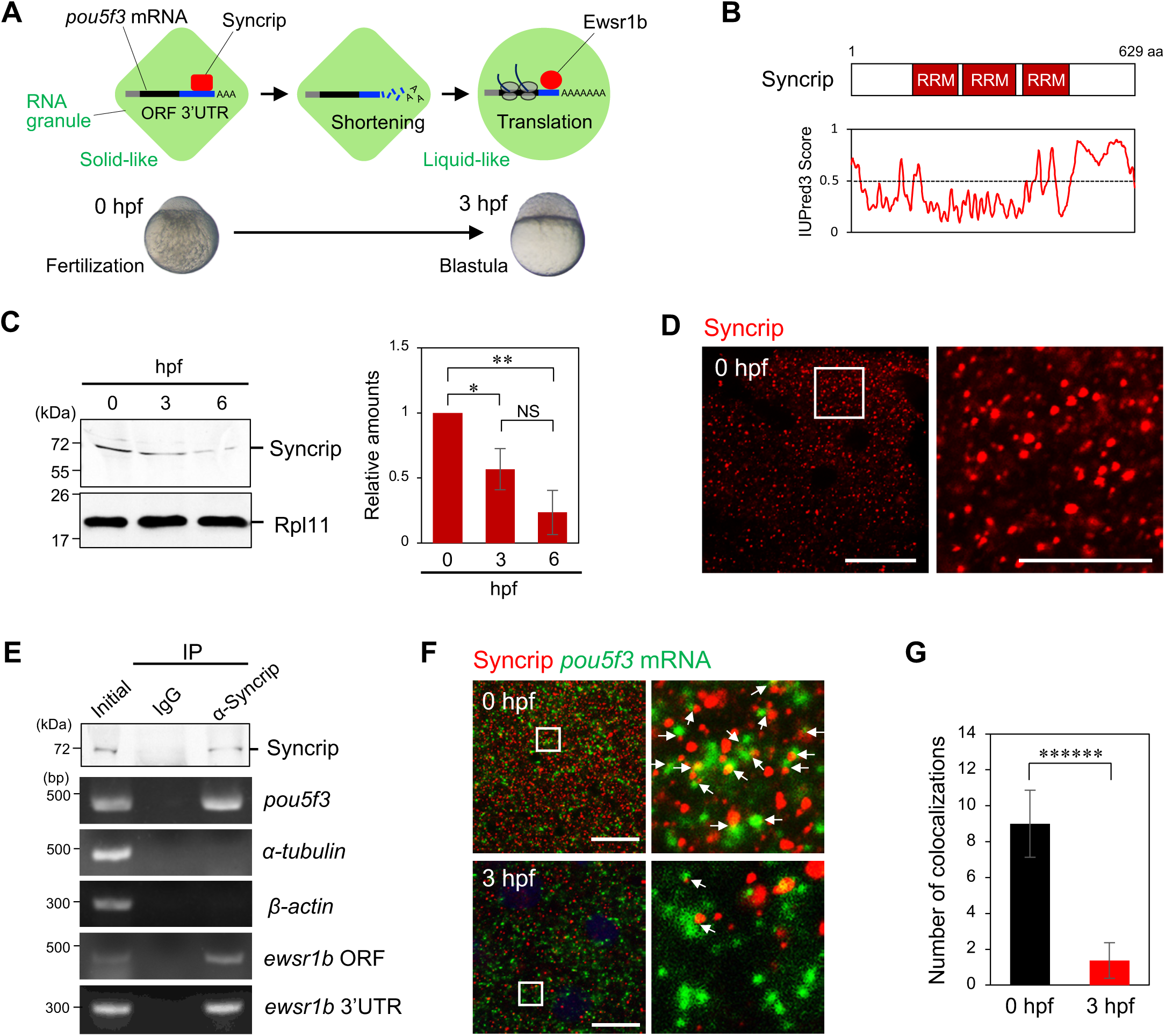
Syncrip is accumulated in eggs, but levels decrease after fertilization. (A) Schematic overview of granule formation, 3′-end shortening, and translation of *pou5f3* mRNA during early development. Syncrip and Ewsr1b were identified as RNA-binding proteins that associate with the long and short 3′UTRs, respectively. (B) Top: Domain structure of zebrafish Syncrip. RRM, RNA recognition motif. Bottom: Predicted intrinsic disorder of Ewsr1b generated by IUPred3. Scores >0.5 indicate regions of disorder. (C) Time-course analysis of Syncrip levels in embryos at various hours post fertilization (hpf). Rpl11 serves as a loading control. Relative Syncrip levels are shown as means ± SD (n = 5). NS, not significant; *p < 0.05; **p < 0.01 (Tukey–Kramer test). (D) Left: Immunofluorescence of Syncrip in embryos at 0 hpf. Scale bar: 20 µm. Right: Enlarged views of the boxed area. Scale bar: 10 µm. (E) Immunoblotting of embryos at 0 hpf before IP (Initial) and after IP using control IgG (IgG) or anti-Syncrip (α-Syncrip) antibody, followed by RT-PCR for *pou5f3*, *α-tubulin*, *β-actin*, and *ewsr1b* mRNAs. Similar results were obtained from two independent experiments. (F) Double staining of *pou5f3* mRNA (green) and Syncrip (red) at 0 and 3 hpf. DNA is shown in blue. Left: High-resolution confocal images. Right: Enlarged views of boxed areas. Arrows indicate colocalized *pou5f3* RNA granules and Syncrip. Scale bars: 20 µm. (G) Quantification of colocalized *pou5f3* RNA granules and Syncrip per 100 µm^2^ at 0 and 3 hpf (means ± SD; n = 8). Similar results were obtained from two independent experiments. ******p < 0.000001 (Student’s *t*-test).

To confirm Syncrip binding to *pou5f3* mRNA in eggs, we performed immunoprecipitation with anti-Syncrip antibody, followed by RT-PCR (IP/RT-PCR). Notably, *pou5f3* mRNA was detected in precipitates from anti-Syncrip antibody but not from control IgG (Figure 1E). Consistently, double staining showed colocalization of Syncrip foci and *pou5f3* RNA granules in embryos at 0 hpf (Figure 1F). This colocalization was reduced in embryos at 3 hpf (Figure 1F and G). Taken together, these results indicate that Syncrip binds to *pou5f3* mRNA in fertilized eggs, and both Syncrip levels and its interaction with mRNA decline over time.

### Syncrip represses *pou5f3* mRNA translation

To assess whether Syncrip regulates *pou5f3* mRNA translation, we expressed GFP-Syncrip by injecting mRNA into fertilized eggs and examined Pou5f3 accumulation. At 6 hpf, GFP-Syncrip localized in cytoplasmic foci, whereas GFP alone diffused throughout cells (Figure 2A). Overexpression of GFP-Syncrip, but not GFP, reduced nuclear accumulation of Pou5f3 (Figure 2A). Immunoblotting confirmed that Pou5f3 levels at 6 hpf were reduced by GFP-Syncrip overexpression but not by GFP alone (Figure 2B). IP/RT-PCR confirmed that GFP-Syncrip interacts with *pou5f3* mRNA (Figure 2C). These results suggest that Syncrip represses *pou5f3* mRNA translation by binding to the mRNA.

**Figure 2.**
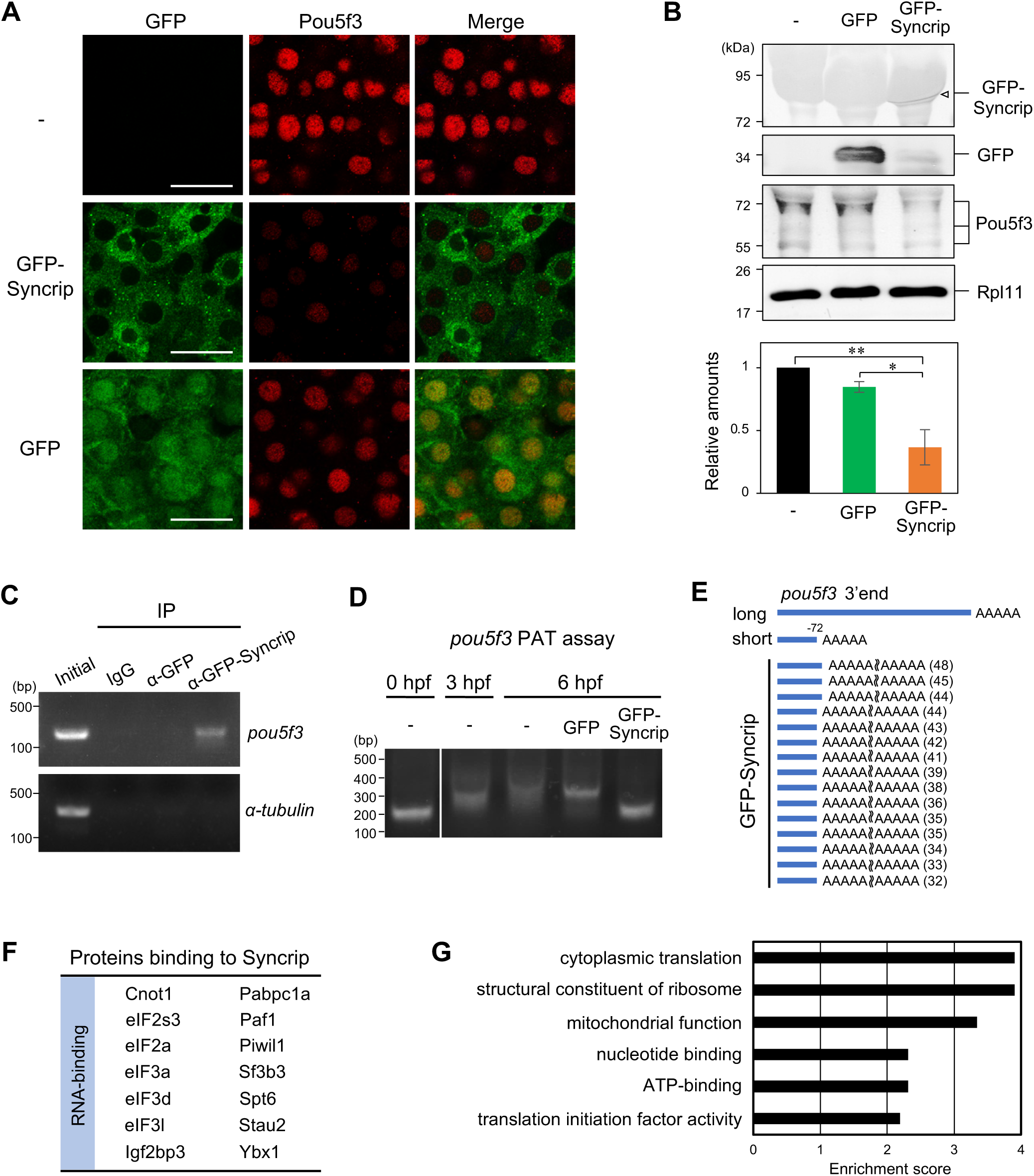
Syncrip represses *pou5f3* mRNA translation. (A) Immunofluorescence of GFP (green) and Pou5f3 (red) in uninjected embryos (-) or embryos injected with GFP-Syncrip (GFP-Syncrip) or GFP (GFP) at 6 hpf. Scale bars: 20 µm. (B) Top: Immunoblotting of GFP-Syncrip, GFP, Pou5f3, and Rpl11 in control and injected embryos at 6 hpf. Bottom: Relative Pou5f3 levels in immunoblotting (means ± SD; n = 3). Similar results were obtained from three independent experiments. *p < 0.05; **p < 0.01 (Tukey–Kramer test). (C) IP/RT-PCR analysis using anti-GFP antibody to detect interactions of GFP or GFP-Syncrip with *pou5f3* and *α-tubulin* mRNAs. Similar results were observed in two independent experiments. (D) PAT assay of *pou5f3* mRNA in embryos at 0 and 3 hpf and in uninjected embryos (-) and those injected with GFP or GFP-Syncrip at 6 hpf. Similar results were obtained from two independent experiments. (E) Sequence analysis of the 3′ ends of *pou5f3* mRNA from GFP-Syncrip-injected embryos at 6 hpf, determined via a PAT assay. Poly(A) tail length are shown, with 3′ ends indicated in blue boxes. (F) Representations of proteins identified via mass spectrometry. (G) GO analysis (molecular function) of proteins interacting with Syncrip.

To investigate the mechanism underlying Syncrip’s repression of *pou5f3* translation, we analyzed *pou5f3* mRNA poly(A) tail length using a poly(A) tail (PAT) assay. Consistent with prior studies [23, 24], the poly(A) tails of *pou5f3* mRNA were elongated at 6 hpf, but this elongation was inhibited by GFP-Syncrip overexpression, although not by GFP alone (Figure 2D). Sequence analysis of PAT assay products showed that 3′-end sequence shortening occurred even in embryos expressing GFP-Syncrip (Figure 2E). These results suggest that Syncrip represses *pou5f3* mRNA translation by inhibiting polyadenylation without blocking 3′-end shortening.

To explore how Syncrip inhibits poly(A) tail elongation, we performed immunoprecipitation of the protein from eggs followed by mass spectrometry. Precipitations with rabbit IgG served as controls. Mass spectrometry analysis identified 87 proteins specifically interacting with Syncrip (Figure 2F and S1C; Table S1). Gene Ontology (GO) analysis revealed enrichment of translational regulators, including Cnot1, a component of the Ccr4–Not complex (Figure 2F and G), which deadenylates poly(A) tails in target mRNAs [39]. These results suggest that Syncrip may repress target mRNA translation by recruiting the Ccr4–Not complex.

### Ewsr1b is synthesized postfertilization and localized in the cytoplasm and nucleus

Ewsr1b was identified as a protein that predominantly binds *pou5f3* mRNA with a shortened 3′UTR [24] (Figure 1A). Ewsr1b is a FET family proteins with conserved domains, including an RNA recognition domain (RRM), zinc finger domain (ZnF), and nuclear localization signal (NLS) (Figure 3A), and contains IDRs across most of its sequence (Figure 3A). In zebrafish, the *ewsr1a* and *ewsr1b* are homologs of human *EWSR1* [40]. We analyzed *ewsr1a* and *ewsr1b* mRNA levels in oocytes and embryos, using RT-PCR and quantitative PCR (qPCR). Results showed that *ewsr1a* mRNA levels were significantly lower than *ewsr1b* mRNA levels in both stages (Figure S2A and B); therefore, we focused on *ewsr1b* for further analysis. The content of *ewsr1b* mRNA remained unchanged up to 6 hpf (Figure S2C), and immunoblotting showed that Ewsr1b protein levels increased from 0 to 6 hpf, especially from 0 to 3 hpf (Figure 3B). Comparing protein levels from 0 to 3 hpf revealed that Ewsr1b content increased at 1 hpf earlier than Pou5f3 content, with the latter increasing at 3 hpf (Figure 3C). Consistently, immunofluorescence showed that Ewsr1b levels increased from 1 to 6 hpf (Figure 3D). Notably, Ewsr1b first localized to cytoplasmic foci and later appeared in the cytoplasm and nucleus from 3 hpf (Figure 3D). Double staining for *pou5f3* mRNA and Ewsr1b protein confirmed that newly synthesized Ewsr1b colocalized with *pou5f3* RNA granules at 3 hpf (Figure 3E and F). These results indicate that Ewsr1b accumulates earlier than Pou5f3 and interacts with *pou5f3* RNA granules.

**Figure 3.**
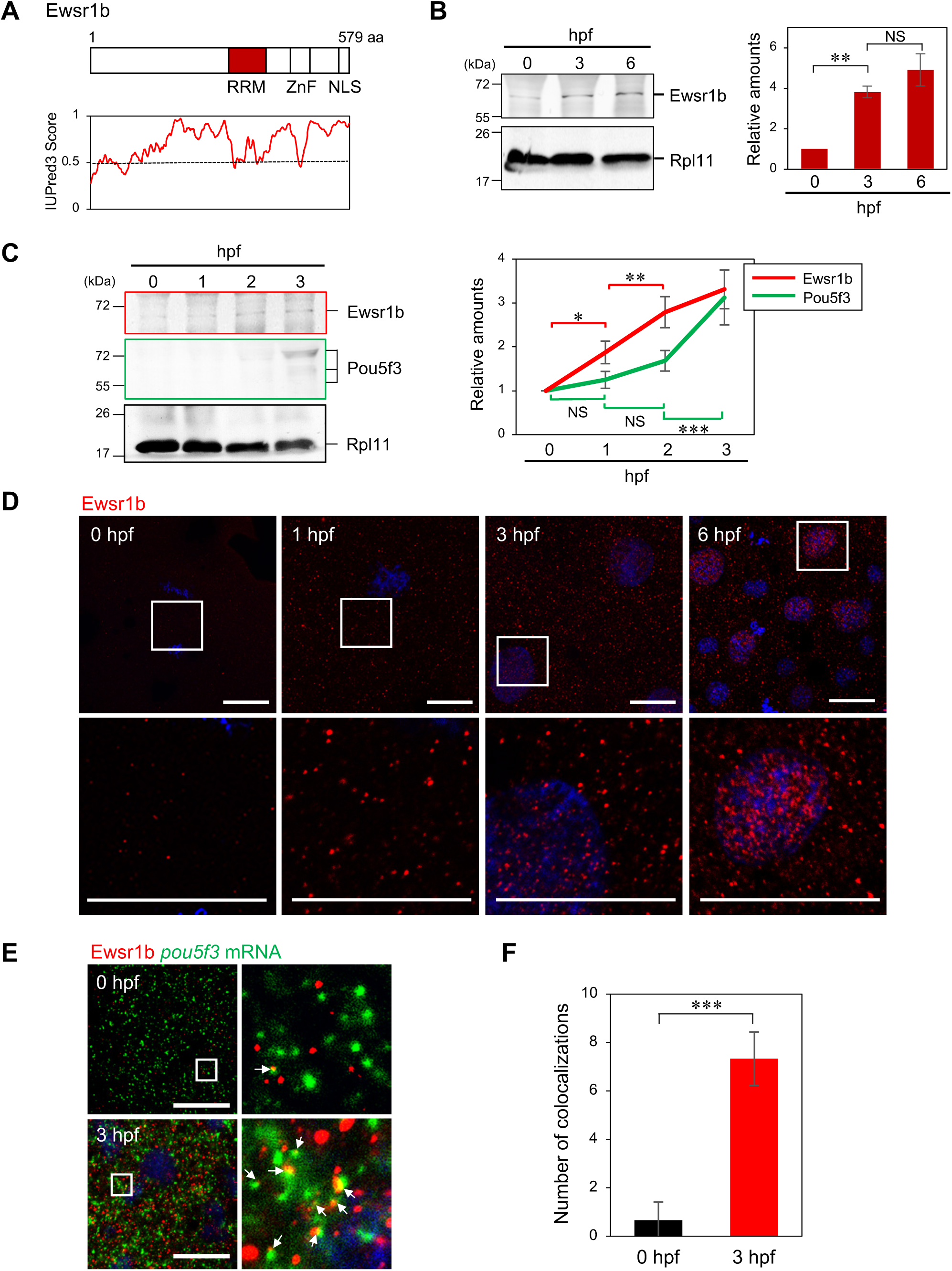
Ewsr1b is synthesized postfertilization and localizes to the cytoplasm and nucleus. (A) Top: Domain structure of zebrafish Ewsr1b. RRM, RNA recognition motif; ZnF, zinc finger domain; NLS, nuclear localization signal. Bottom: Predictions of intrinsic disorder in Ewsr1b via IUPred3. (B) Left: Time-course analysis of Ewsr1b levels in embryos at 0, 3, and 6 hpf. Right: Relative protein amounts (means ± SD; n = 3). NS, not significant; **p < 0.01 (Tukey–Kramer test). (C) Left: Time-course analysis of Ewsr1b and Pou5f3 protein levels in embryos at 0, 1, 2, and 3 hpf. Right: Relative Ewsr1b and Pou5f3 levels at 0, 1, 2, and 3 hpf (means ± SD; n = 3). NS, not significant; *p < 0.05; **p < 0.01; ***p < 0.001 (Tukey–Kramer test). (D) Immunofluorescence of Ewsr1b at 0, 1, 3, and 6 hpf. DNA is shown in blue. Upper panels: High-resolution confocal images; lower panels: enlarged views of boxed regions. Similar results were obtained from three independent experiments. Scale bars: 20 µm. (E) Double staining of *pou5f3* mRNA (green) and Ewsr1b (red) at 0 and 3 hpf. DNA is shown in blue. Left: High-resolution confocal images; right: enlarged views of boxed regions. Arrows indicate colocalized *pou5f3* mRNA and Ewsr1b. Scale bars: 20 µm. (F) Quantification of colocalized *pou5f3* mRNA and Ewsr1b signals per 100 µm^2^ at 0 and 3 hpf (means ± SD; n = 3). Similar results were obtained from three independent experiments. *****p < 0.00001 (Student’s *t*-test).

### Ewsr1b promotes *pou5f3* mRNA translation

To determine whether Ewsr1b regulates the translation of *pou5f3* mRNA, we performed triple staining of *pou5f3* mRNA, Ewsr1b, and Pou5f3 in embryos at 3 hpf. Pou5f3 was detected at the sites where Ewsr1b colocalized with *pou5f3* RNA granules, and Ewsr1b was present at sites where *pou5f3* RNA granules and Pou5f3 were colocalized (Figure 4A and B), suggesting a role for Ewsr1b in the translational activation of *pou5f3* mRNA. Ewsr1b also colocalized with Pou5f3 in the nucleus as well as the cytoplasm (Figure 4C). A proximity ligation assay (PLA), enabling the detection of two molecules existing in proximity (<40 nm), confirmed interactions between Ewsr1b and Pou5f3, with signals detected in the cytoplasm at 2 hpf and predominantly in the nucleus by 3 hpf (Figure S3A and B), confirming the Ewsr1b–Pou5f3 interaction in both compartments.

**Figure 4.**
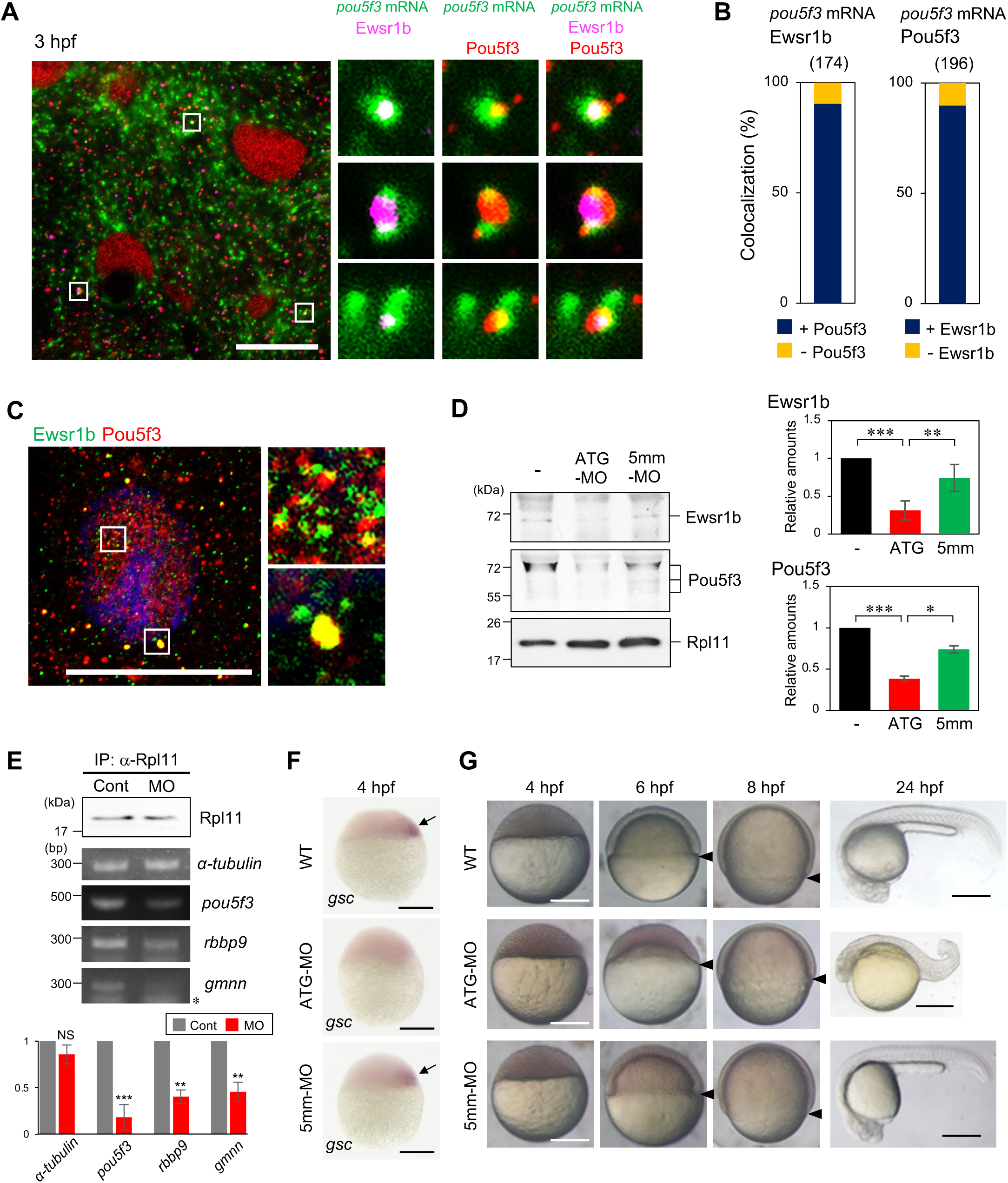
Ewsr1b promotes *pou5f3* mRNA translation. (A) Triple staining of *pou5f3* mRNA (green), Ewsr1b (magenta), and Pou5f3 (red) in embryos at 3 hpf. Left: High-resolution confocal image; right: enlarged views of boxed regions. Scale bar: 20 µm. (B) Left: Percentage of colocalizations with (+) or without (−) Pou5f3 at sites where *pou5f3* mRNA and Ewsr1b are colocalized. Right: Percentage of colocalizations with (+) or without (−) Ewsr1b at sites where *pou5f3* mRNA and Pou5f3 are colocalized. (C) Immunofluorescence of Ewsr1b (green) and Pou5f3 (red) at 3 hpf. DNA is shown in blue. Left: High-resolution confocal image; right: enlarged views of boxed area. Similar results were obtained from three independent experiments. Scale bar: 20 µm. (D) Left: Immunoblotting of Ewsr1b, Pou5f3, and Rpl11 in uninjected embryos (−) and those injected with *ewsr1b*-ATG-MO (ATG-MO) or *ewsr1b*-5mm-MO (5mm-MO) at 6 hpf. Right: Quantification of immunoblotting (means ± SD; n = 3). *p < 0.05; **p < 0.01; ***p < 0.001 (Tukey–Kramer test). (E) Upper: Immunoblotting of uninjected embryos (Cont) and those injected with ATG-MO (MO) at 4 hpf after IP with anti-Rpl11 antibody and RT-PCR for *α-tubulin*, *pou5f3*, *rbbp9*, and *gmnn* mRNAs. An asterisk indicates nonspecific bands. Lower panel: RT-qPCR of *pou5f3*, *α-tubulin*, *rbbp9*, and *gmnn* mRNAs in uninjected embryos (Cont) and those injected with ATG-MO (MO) at 4 hpf after IP with anti-Rpl11 antibody (means ± SD; n = 3). Similar results were obtained from three independent experiments. NS, not significant; **p < 0.01; ***p < 0.001 (Student’s *t*-test). (F) Whole-mount *in situ* hybridization of *gsc* mRNA in wild-type embryos and those injected with ATG-MO and 5mm-MO at 4 hpf. Arrows indicate zygotic *gsc* transcripts. Similar results were obtained from two independent experiments. (G) Lateral views of wild-type embryos and those injected with ATG-MO and 5mm-MO at 4, 6, 8, and 24 hpf. Scale bars: 300 µm. Similar results were obtained from six independent experiments.

To investigate the function of Ewsr1b, we knocked down its expression using a morpholino oligonucleotide (MO) targeting the start codon of *ewsr1b* mRNA (ATG-MO; Figure S4A), with a control MO containing 5-nucleotide mismatches (5mm-MO) used for comparison. Injection of ATG-MO, but not 5mm-MO, inhibited Ewsr1b synthesis (Figure 4D) and led to reduced Pou5f3 levels in Ewsr1b-deficient embryos (Figure 4D). To assess the effects of Ewsr1b on *pou5f3* mRNA translation, we immunoprecipitated the 60S ribosome large subunit protein Rpl11 and detected translated mRNAs via RT-PCR (Rpl11-IP/RT-PCR). Results showed that *pou5f3* mRNA was translated at 4 hpf and that translation was reduced in Ewsr1b-deficient embryos (Figure 4E). We also tested whether Ewsr1b affects the physical state of *pou5f3* RNA granules by treating embryos with hexanediol, which dissolves liquid-phase droplets. In controls, the number of *pou5f3* RNA granules decreased at 3 hpf, whereas in Ewsr1b-deficient embryos, the number remain unchanged (Figure S4B), indicating that Ewsr1b promotes the transition of *pou5f3* RNA granules from a solid- to liquid-like state, thereby facilitating translation. Consistent with the Pou5f3 reduction, Ewsr1b-deficient embryos failed to activate zygotic gene transcription at the initiation of ZGA (Figure 4F and S4C), indicating that Ewsr1b-mediated translational activation is crucial for ZGA.

Ewsr1b knockdown via ATG-MO injection, but not control 5mm-MO, caused severe developmental defects, including delayed epiboly from 6 hpf and overall shrinkage of the embryo by 24 hpf (Figure 4G and S4D). Similar defects were observed with a second MO targeting the 5′UTR of *ewsr1b* mRNA (Figure S4A, E and F), and these were rescued via coinjection of GFP-*ewsr1b* mRNA lacking the 5′UTR (Figure S4E and F). Taken together, these findings demonstrate that Ewsr1b is crucial for early embryonic development.

### Two *ewsr1b* mRNA variants are translated at distinct times

As *ewsr1b* mRNA is translated before the translational activation of *pou5f3* mRNA, we investigated how *ewsr1b* is translationally regulated. Unexpectedly, the PAT assay revealed that one *ewsr1b* transcript possessed a short 3′UTR comprising only 16 nucleotides (Short-3′UTR; Figure 5A and B). Its poly(A) tail was rapidly elongated between 0 and 1 hpf (Figure 5B). The PAT assay was also performed using primers for database sequences of *ewsr1b* (Figure 5A). Results revealed a second *ewsr1b* transcript in eggs with a long 3′UTR comprising 302 nucleotides (Long-3′UTR; Figure 5A and B), exhibiting gradual poly(A) tail elongation from 0 to 3 hpf (Figure 5B). Both variants were also identified in 3′-end RNA sequencing [24, 41] (Figure 5A). Hereafter, these variants are referred as *ewsr1b*-3′Short and *ewsr1b*-3′Long mRNAs.

**Figure 5.**
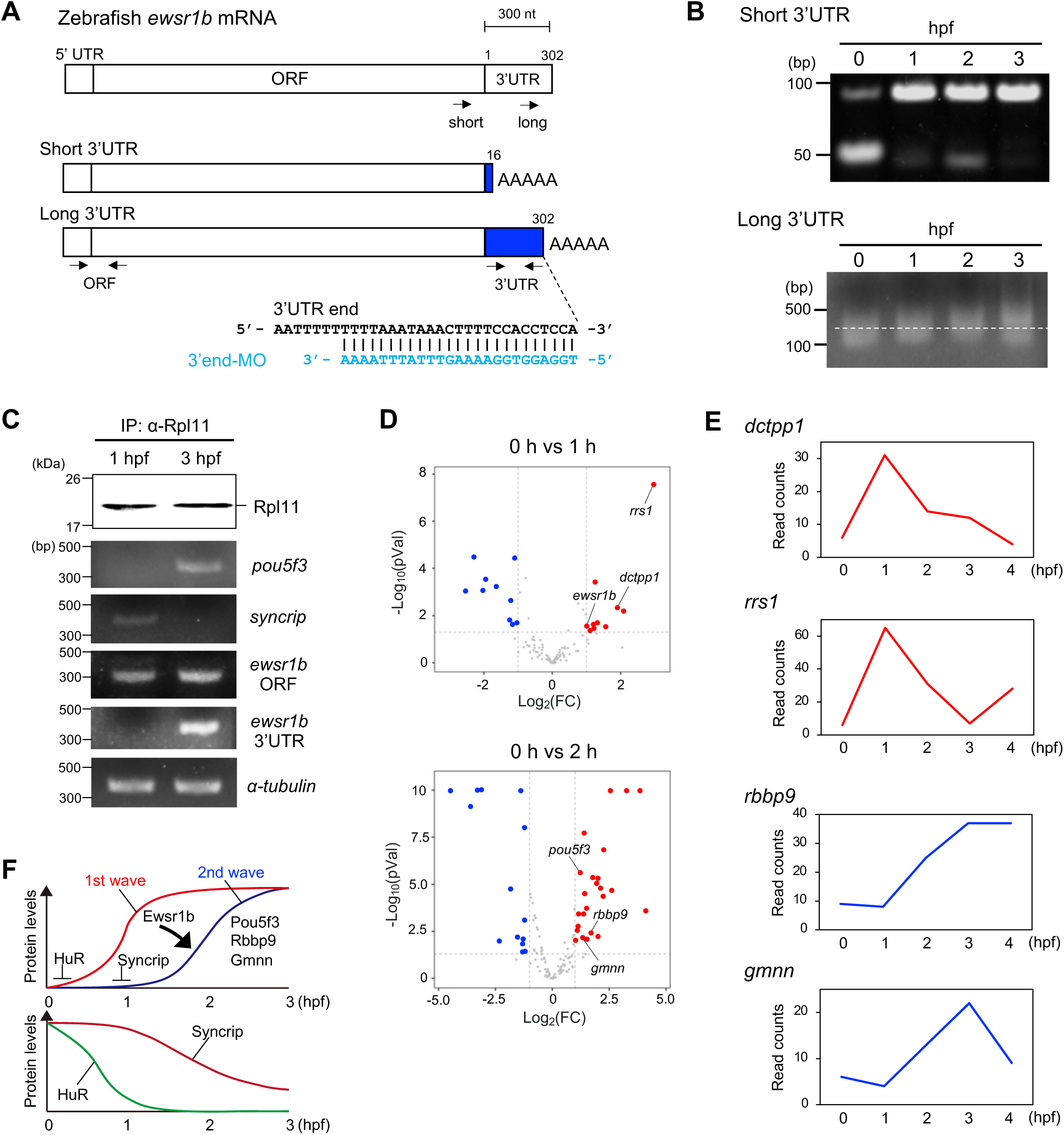
Two *ewsr1b* mRNA variants are translated at distinct times. (A) Top: Schematic of full-length *ewsr1b* mRNA. ORF, open reading frame. Arrows indicate forward primers for PAT assays of short and long 3′UTRs, respectively. Middle: Schematics for *ewsr1b* mRNAs harboring short (Short 3′UTR) and long (Long 3′UTR) 3′UTRs. Arrows indicate primers used to detect the *ewsr1b* ORF and long 3′UTR. Bottom: Sequences of *ewsr1b* mRNA 3′ end targeted by *ewsr1b*-3′end-MO (3′end-MO). (B) PAT assays of *ewsr1b* mRNA harboring short (Short 3′UTR) or long (Long 3′UTR) 3′UTR in embryos at 0, 1, 2, and 3 hpf. Similar results were obtained from three independent experiments. (C) Top: Immunoblotting of embryos at 1 and 3 hpf after IP with anti-Rpl11 (α-Rpl11) antibody. Bottom: RT-PCR for *pou5f3*, *syncrip*, *ewsr1b* mRNA ORF (*ewsr1b* ORF), *ewsr1b* mRNA 3′UTR (*ewsr1b* 3′UTR), and *α-tubulin* mRNA. Similar results were obtained from two independent experiments. (D) Volcano plots showing differential transcript abundance from RNA sequencing between 0 and 1 hpf (top) and between 0 and 2 hpf (bottom). (E) Read count trajectories of *dctpp1*, *rrs1*, *rbbp9*, and *gmnn* mRNAs from 0 to 4 hpf. (F) Schematics of translational activation waves: first wave at ∼1 hpf and second at ∼2 hpf. Syncrip represses the translation of second-wave transcripts: HuR represses first-wave transcripts.

To compare translational timings for *ewsr1b*-3′Short and *ewsr1b*-3′Long mRNAs, we performed Rpl11-IP/RT-PCR in embryos at 1 and 3 hpf. As expected, *pou5f3* mRNA and *syncrip* mRNA were only detected in embryos at 3 hpf and 1 hpf, respectively (Figure 5C). In contrast, using primers in the *ewsr1b* open reading frame (ORF), *ewsr1b* was detected at 1 and 3 hpf (Figure 5A and C). However, *ewsr1b*-3′Long mRNA was only detected at 3 hpf (Figure 5A and C), indicating that *ewsr1b*-3′Short mRNA is translated earlier. Because the later-stage timing of *ewsr1b*-3′Long mRNA translation overlaps with that of *pou5f3* mRNA (Figure 5C), we examined whether Syncrip represses the *ewsr1b*-3′Long mRNA translation. Immunoprecipitation followed by RT-PCR and 3′ rapid amplification of cDNA ends (3′RACE) showed that Syncrip binds *ewsr1b*-3′Long, but not *ewsr1b*-3′Short, mRNA in eggs (Figure 1E and S5B). In addition, Syncrip overexpression reduced Ewsr1b accumulation at 6 hpf (Figure S5C). Thus, Syncrip appears to repress *ewsr1b*-3′Long mRNA translation in a similar manner to *pou5f3* translation.

To assess broader translational dynamics, we conducted immunoprecipitation of Rpl11 followed by RNA sequencing in short intervals from 0 to 4 hpf. This analysis identified 10 mRNAs with >2-fold increased translation from 0 to 1 hpf (Figure 5D and E; Table S2). Additionally, >20 mRNAs showed elevated translation after 2 hpf (Figure 5D and E; Table S2). Temporal changes in *ewsr1b* and *pou5f3* mRNA translation were consistent with immunoblotting, PAT assay, and RT-PCR results (Figure S5D). To determine whether Ewsr1b influences the translational activation of mRNAs after 2 hpf, we examined Ewsr1b knockdown effects on *rbbp9* and *gmnn* mRNA translation. Similar to *pou5f3* mRNA, these mRNAs showed significantly decreased translation in Ewsr1b-deficient embryos (Figure 4E), suggesting that Ewsr1b synthesized at 0–1 hpf induces the activation of specific mRNAs from 2 hpf. Overall, these results indicate that temporal translation of mRNAs is coordinated in embryos through sequential translation waves, with the first inducing the second (Figure 5F). Similar to *ewsr1b*-3′Short mRNA, some mRNAs translated in the first wave harbored short 3′UTRs (Figure S5A).

### Difference in 3′UTR length of *ewsr1b* mRNA determines localization and function of synthesized Ewsr1b

Ewsr1b initially localized in the cytoplasm and later accumulated in both the cytoplasm and nucleus (Figure 3D). To assess whether translation of the two *ewsr1b* mRNA variants contributes to Ewsr1b localization, we examined the distribution of Ewsr1b synthesized from reporter mRNAs containing either the short or long 3′UTR variant (Figure 6A). Notably, GFP-Ewsr1b translated from the Short-3′UTR mRNA localized in the cytoplasm, whereas GFP-Ewsr1b translated from the Long-3′UTR mRNA localized in the nucleus (Figure 6B).

**Figure 6.**
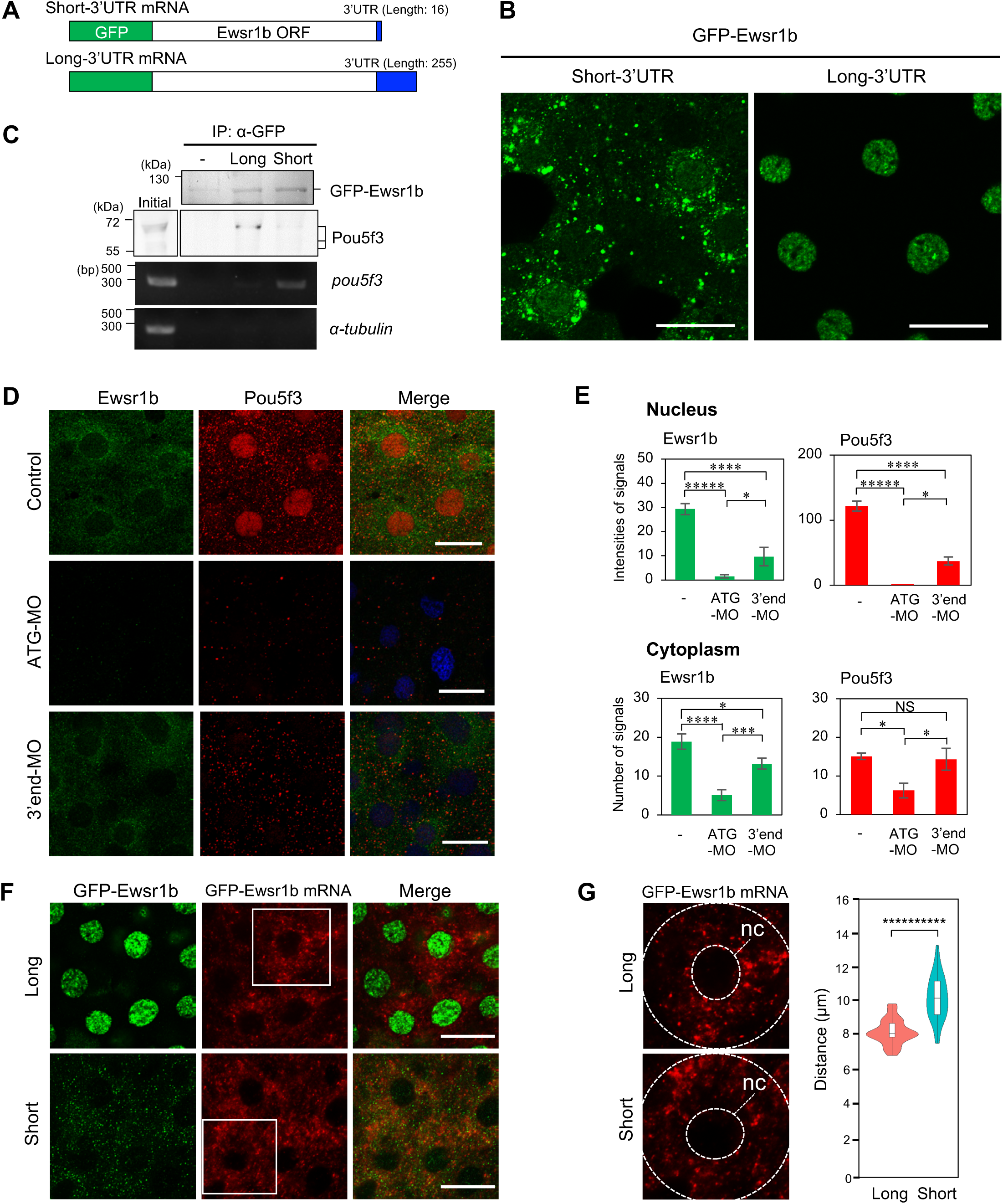
Difference in 3′UTR length of *ewsr1b* mRNA determines Ewsr1b localization and function. (A) Schematics of GFP-Ewsr1b mRNA carrying Short-3′UTR or Long-3′UTR. (B) Localization of GFP-Ewsr1b synthesized from GFP-Ewsr1b mRNA carrying Short-3′UTR or Long-3′UTR in embryos at 6 hpf. Similar results were obtained from three independent experiments. Scale bars: 20 µm. (C) Coimmunoprecipitation and IP/RT-PCR analyses of GFP-Ewsr1b synthesized from GFP-Ewsr1b mRNA carrying Long-3′UTR or Short-3′UTR. Upper panel: Immunoblotting of uninjected embryos (−) and those injected with GFP-Ewsr1b mRNA carrying Long-3′UTR (Long) or Short-3′UTR (Short) at 6 hpf before (Initial) and after (α-GFP) immunoprecipitation with anti-GFP antibody. Lower panel: RT-PCR for *pou5f3* and *α-tubulin* mRNAs. Similar results were obtained from two independent experiments. (D) Immunofluorescence of Ewsr1b (green) and Pou5f3 (red) in uninjected embryos (Control) and those injected with *ewsr1b*-ATG-MO (ATG-MO) and *ewsr1b*-3′end-MO (3′end-MO) at 4 hpf. Scale bars: 20 µm. (E) Upper panel: Quantification of nuclear signal intensities for Ewsr1b and Pou5f3 (means ± SD; n = 18). Lower panel: Number of cytoplasmic signals for Ewsr1b and Pou5f3 per 100 µm^2^ (means ± SD; n = 18). NS, not significant; *****p < 0.00001; ****: p < 0.0001; ***p < 0.001; *p < 0.05 (Tukey–Kramer test). Similar results were obtained from two independent experiments. (F) Double staining of GFP-Ewsr1b (green) and GFP-Ewsr1b mRNA (red) in embryos injected with GFP-Ewsr1b mRNA carrying Long-3′UTR (Long) or Short-3′UTR (Short) at 4 hpf. Scale bars: 20 µm. (G) Violin plots showing distances from the nuclear center to signals of GFP-Ewsr1b mRNA carrying Long-3′UTR or Short-3′UTR (means ± SD; n = 80). Similar results were obtained from two independent experiments. **********p < 0.0000000001 (Student’s *t*-test).

To further characterize this differential localization, we immunoprecipitated GFP-Ewsr1b and assessed its interaction with Pou5f3 protein and *pou5f3* mRNA via immunoblotting and RT-PCR, respectively. Immunoblotting revealed no difference in molecular weight between GFP-Ewsr1b synthesized from either 3′UTR variant (Figure 6C). However, Pou5f3 was coprecipitated only with GFP-Ewsr1b synthesized from Long-3′UTR mRNA (Figure 6C), whereas *pou5f3* mRNA coprecipitated only with GFP-Ewsr1b from Short-3′UTR mRNA (Figure 6C). Thus, Ewsr1b translated from Long-3′UTR mRNA localizes to the nucleus and interacts with Pou5f3, whereas Ewsr1b from Short-3′UTR mRNA localizes to the cytoplasm and binds *pou5f3* mRNA. Furthermore, the proximal 90 nucleotides of the *ewsr1b* 3′UTR were sufficient to promote nuclear localization of Ewsr1b (Figure S6A and B), suggesting that specific 3′UTR sequences, rather than overall length, determine protein localization.

To investigate the functional differences in Ewsr1b synthesized from each variant, we inhibited the translation of *ewsr1b*-3′Long mRNA using a MO targeting its 3′ end (3′end-MO; Figure 5A). Immunofluorescence showed that blocking Ewsr1b translation from both mRNAs using ATG-MO reduced Ewsr1b and Pou5f3 protein levels in the nucleus and cytoplasm (Figure 6D and E). In contrast, inhibition of *ewsr1b*-3′Long mRNA translation alone specifically decreased Ewsr1b and Pou5f3 levels in the nucleus (Figure 6D and E), with Ewsr1b content only slightly decreased in the cytoplasm, and Pou5f3 content remaining unchanged (Figure 6D and E). These results suggest that Ewsr1b translated from *ewsr1b*-3′Long mRNA accumulates in the nucleus and stabilizes Pou5f3 protein expression, whereas Ewsr1b synthesized from *ewsr1b*-3′Short mRNA functions in the cytoplasm and promotes mRNA translation. Inhibition of the long variant did not affect early development until the gastrulation stage but led to aberrant phenotypes, such as small body size and a curled tail during tailbud stages (Figure S7A and B), highlighting the importance of *ewsr1b*-3′Long mRNA translation for late-stage developmental processes.

### Distributions of mRNA influence the subcellular localization of synthesized Ewsr1b

To determine whether mRNA localization correlates with protein localization, we analyzed the distribution of reporter mRNAs using fluorescence in situ hybridization (FISH) with a probe targeting *gfp* (Figure 6F). Long-3′UTR mRNA localized to the perinuclear region, whereas Short-3′UTR mRNA localized to the cell’s outer region (Figure 6G). To assess the distribution of endogenous *ewsr1b* mRNAs, we performed FISH using separate probes targeting the ORF (green) and 3′UTR (red) regions, respectively (Figure S8A), distinguishing *ewsr1b* mRNAs harboring long and short 3′UTRs (Figure S8B). Measuring the distance from the nucleus’ center to the signal (Figure S8C), we found that *ewsr1b*-3′Short mRNA localized in the peripheral cytoplasm, whereas ewsr1b-3′Long mRNA localized nearer the nucleus (Figure S8C). These findings indicate that 3′UTR length governs mRNA localization, which in turn affects the translated Ewsr1b protein’s subcellular distribution.

We next examined the mechanism underlying nuclear accumulation of Ewsr1b. The protein contains a NLS, typically recognized by nuclear transport proteins, such as importins [42, 43]. We hypothesized that if an importin family protein bound to *ewsr1b*-3′Long mRNA but not *ewsr1b*-3′Short mRNA, Ewsr1b synthesized from *ewsr1b*-3′Long mRNA would be effectively transported into the nucleus. Based on a report of human EWSR1 binding to Importin β1 [44], we tested whether Importin β1 interacts with *ewsr1b* mRNA. Immunofluorescence showed that Importin β1 localized to both the nucleus and cytoplasm (Figure 7A). IP/RT-PCR revealed that Importin β1 binds to *ewsr1b* mRNA (Figure 7B), and double staining demonstrated colocalization with the 3′UTR of *ewsr1b*-3′Long mRNA (Figure 7C). PLA analysis confirmed interactions between Importin β1 and Ewsr1b protein in both compartments at 3 hpf, resulting in nuclear accumulation by 6 hpf (Figure S9A).

**Figure 7.**
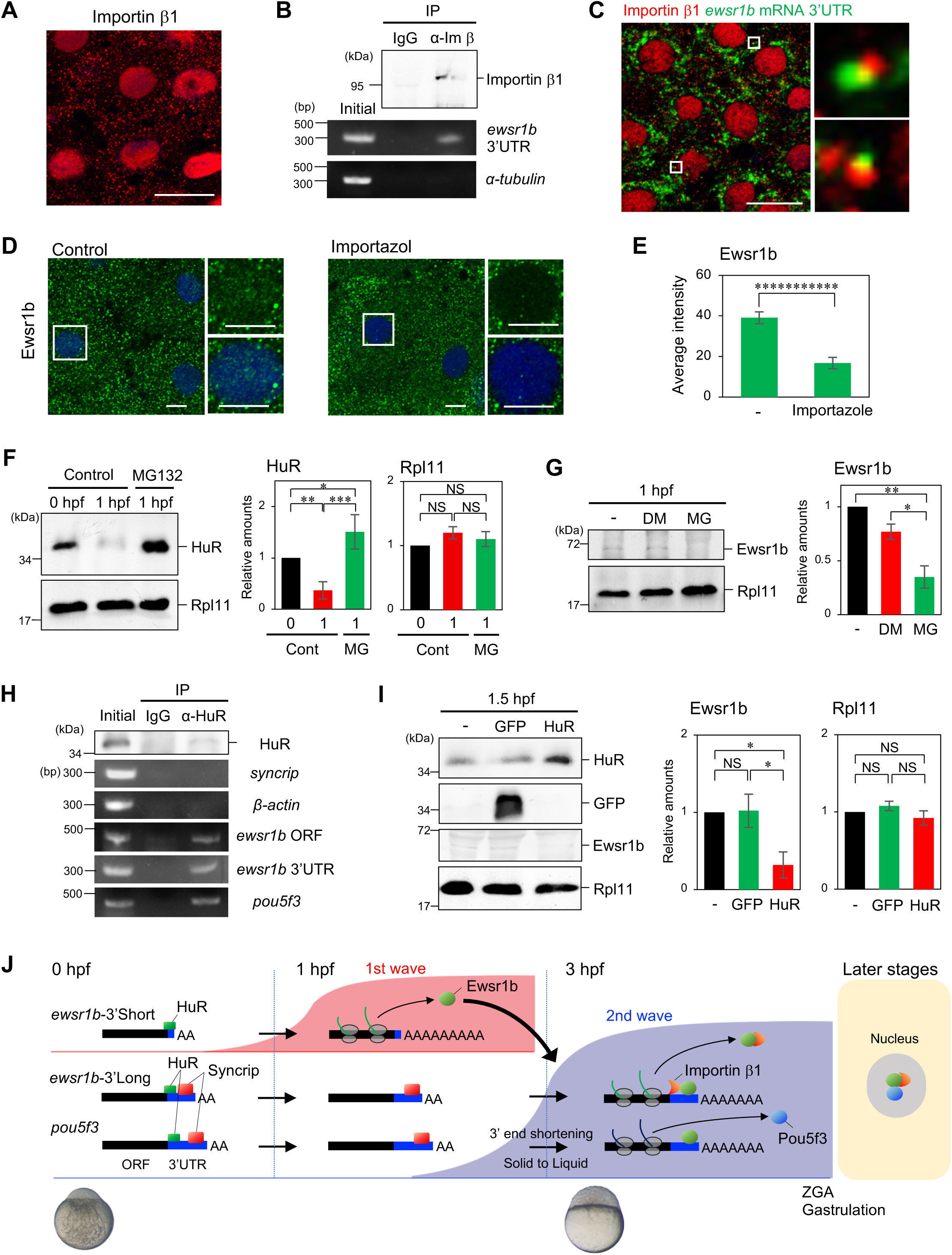
Regulation of *ewsr1b*-3′Long and *ewsr1b* -3′Short mRNAs. (A) Immunofluorescence of Importin β1 in embryos at 3 hpf. (B) Immunoblotting of embryos at 3 hpf following IP with control IgG (IgG) or anti-Importin β1 (α-Im β) antibody, and RT-PCR for *ewsr1b*-3′Long and *α-tubulin* mRNAs. Similar results were obtained from two independent experiments. (C) Double staining of Importin β1 (red) and the *ewsr1b*-3′Long mRNA 3′UTR (green) in embryos at 3 hpf. Left: High-resolution confocal image; right: enlarged views of the boxed area. Similar results were obtained from two independent experiments. (D) Immunofluorescence of Ewsr1b in uninjected embryos (Control) or embryos injected with Importazole at 3 hpf. DNA is shown in blue. Enlarged views of the boxed area with or without DNA staining are shown on the right side. Scale bars, 10 µm. (E) Quantification of average signal intensity in the nucleus per 25 µm^2^. (means ± SD; n = 10). ***********p < 0.00000000001 (Student’s *t*-test). (F) Left: Immnunoblotting of HuR and Rpl11 in uninjected embryos (Control) at 0 and 1 hpf and embryos injected with MG132 at 1 hpf. Middle and Right: Quantification of immunoblotting (means ± SD; n = 3). NS, not significant; *p < 0.05; **p < 0.01; ***p < 0.001 (Tukey–Kramer test). (G) Left: Immunoblotting of Ewsr1b and Rpl11 in uninjected embryos (−) and those injected with DMSO or MG132 at 1 hpf. Right: Quantification of immunoblotting (means ± SD; n = 3). Similar results were obtained from three independent experiments. *p < 0.05; **p < 0.01 (Tukey–Kramer test). (H) Immunoblotting of embryos at 0 hpf before IP (Initial) and after IP using control IgG (IgG) or anti-HuR (α-HuR) antibody, followed by RT-PCR for *syncrip*, *β-actin*, *ewsr1b*, and *pou5f3* mRNAs. Similar results were obtained from two independent experiments. (I) Left: Immunoblotting of HuR, GFP, Ewsr1b, and Rpl11 in uninjected embryos (-) or embryos injected with GFP or HuR at 1.5 hpf. Right: Relative Ewsr1b and Rpl11 levels in immunoblotting (means ± SD; n = 3). Similar results were obtained from three independent experiments. NS, not significant; *p < 0.05 (Tukey–Kramer test). (J) Schematic summary of the translational and cotranslational control of *ewsr1b*-3′Short, *ewsr1b*-3′Long, and *pou5f3* mRNAs in embryos at 0, 1 and 3 hpf and in later developmental stages.

To assess whether Importin β1 transports Ewsr1b in the nucleus, we treated embryos with an Importin β-specific inhibitor importazole [45]. This treatment prevented Importin β1 accumulation in the nucleus (Figure S9B and C). Notably, nuclear localization of endogenous Ewsr1b and GFP-Ewsr1b translated from the Long-3′UTR mRNA were significantly perturbed (Figure 7D and E; Figure S9D and E). These results support a model in which 3′UTR sequences of *ewsr1b*-3′Long mRNA recruit Importin β1 to facilitate nuclear import of Ewsr1b.

### First wave of *ewsr1b* mRNA translation depends on proteasome activity

To explore how *ewsr1b*-3′Short mRNA is rapidly translated after fertilization, we investigated the role of proteasome-mediated protein degradation, which is known to occur immediately after fertilization in fish and mammals [46, 47]. We hypothesized that such degradation removes translational repressors of *ewsr1b*-3′Short mRNA. In support of this, we found that an RNA-binding protein HuR (also known as Elavl1) was rapidly degraded within 1 h after fertilization (Figure 7F). This degradation was dependent on proteasome activity because treatment with the proteasome inhibitor MG132 prevented HuR reduction (Figure 7F). Furthermore, MG132 suppressed Ewsr1b synthesis at 1 hpf (Figure 7G and S10A), despite almost normal mitotic cleavage (Figure S10B). A PAT assay and Rpl11-IP/RT-PCR demonstrated that MG132 inhibited poly(A) tail elongation and translation of *ewsr1b*-3′Short mRNA at 1 hpf (Figure S10C and D). Thus, proteolysis postfertilization appears necessary for translational activation of *ewsr1b*-3′Short mRNA.

To explore the relationship between HuR degradation and *ewsr1b* mRNA translation, we immunoprecipitated HuR in eggs and examined their interaction by RT-PCR and 3′RACE. Notably, *syncrip* and *β-actin* mRNAs were not detected in HuR precipitations, whereas *ewsr1b* and *pou5f3* mRNAs were detected (Figure 7H). 3′RACE revealed that both *ewsr1b*-3′Short and −3′Long mRNAs interacted with HuR (Figure S10E). We then analyzed the effect of HuR-overexpression by injecting mRNA into fertilized eggs. Overexpression of HuR, but not GFP, prevented *ewsr1b* mRNA translation (Figure 7I). Taken together, these results suggest that HuR represses *ewsr1b*-3′Short mRNA translation, which is derepressed by proteasome-dependent HuR degradation, initiating the first translation wave.

### Ewsr1b synthesized in the first translation wave exhibits liquid-like properties

Finally, we characterized the biophysical properties of Ewsr1b synthesized from ewsr1b-3′Short mRNA. Time-lapse imaging of GFP-Ewsr1b revealed droplet fusion, a property of liquid-phase behavior (Figure S11A). Furthermore, fluorescence recovery after photobleaching (FRAP) analysis demonstrated rapid fluorescence recovery following photobleaching (Figure S11B and C), consistent with a dynamic liquid-like state. Given that Ewsr1b colocalizes with *pou5f3* RNA granules (Figure 4A and B) and is required for their transition from a solid- to liquid-like state (Figure S4B), these results support a model in which Ewsr1b promotes the biophysical transformation of RNA granules into liquid droplets upon binding.

## DISCUSSION

Precise regulation of protein syntheses is essential for the progression of nearly all biological processes. However, the molecular and cellular systems that coordinate translation in temporal and spatial dimensions, particularly during prolonged and multilayered processes remain largely unclear. In this study, we identified sequential waves of translational activation of dormant maternal mRNAs following fertilization and revealed previously unknown molecular principles for coordinating mRNA translation, protein function, and developmental processes (Figure 7J). The first translation wave is initiated immediately after fertilization (Figures 3 and 5) and involves mRNAs harboring short 3′UTRs. Ewsr1b translated from *ewsr1b*-3′Short mRNA localizes to the cytoplasm and activates the second translation wave, including *pou5f3* mRNA translation (Figures 3 and 4). Ewsr1b is also synthesized during the second wave from mRNAs harboring a long 3′UTR (Figure 5). Importin β1 binds to *ewsr1b*-3′Long mRNA and facilitate nuclear transport of the newly synthesized Ewsr1b (Figure 7). In the nucleus, Ewsr1b colocalizes with Pou5f3 contributing to its nuclear maintenance and the regulation of later developmental processes (Figure 6). In contrast, the translation of *ewsr1b*-3′Short mRNA is repressed by HuR (Figure 7). Syncrip binds to *pou5f3* and *ewsr1b*-3′Long mRNAs and represses their translation until the second wave (Figures 1 and 2). Although HuR binds all three mRNAs (Figure 7H and S10E), HuR degradation triggers *ewsr1b*-3′Short mRNA translation but not others because of the absence of Syncrip on *ewsr1b*-3′Short mRNA (Figure 1E and S5B). Ewsr1b-mediated translational activation is crucial for embryonic development including ZGA and gastrulation (Figure 4).

### Translational repression of dormant mRNAs by Syncrip in early development

Originally identified in mouse neural tissues, Syncrip has been implicated in a range of biological processes, particularly in neuronal development through posttranscriptional regulation [25, 29, 30, 48–50]. However, its role in early embryogenesis and its interacting partners had remained unclear. We showed that Syncrip represses the translation of *pou5f3* and *ewsr1b*-3′Long mRNAs during early development (Figures 1, 2, and S5C). Given that numerous mRNAs were translated during the second wave (Figure 5) and that abundant many Syncrip foci were present in the egg cytoplasm (Figure 1), Syncrip may regulate the translation of these mRNAs similarly to *pou5f3*. Furethermore, our mass spectrometry analysis revealed that Syncrip interacts with multiple RNA-binding proteins involved in mRNA regulation, including Cnot1 and Igf2bp3 (Figure 2F and G). Cnot1 is a core component of the Ccr4–Not complex, which mediates mRNA deadenylation and degradation in eukaryotic cells [39], whereas Igf2bp3 has been shown to stabilize maternal mRNAs in early-stage zebrafish embryos [51]. These findings suggest that Syncrip acts as a hub for RNA regulatory proteins that govern maternal mRNA translation during early development. Furthermore, overexpression of GFP-Syncrip suppressed poly(A) tail elongation in *pou5f3* mRNA after the 3′ ends had been shortened (Figure 2D and E), consistent with prior studies indicating that cytoplasmic polyadenylation is important for translational activation of dormant mRNAs postfertilization [24, 52].

### Translational activation of dormant mRNAs by Ewsr1b during early development

Ewsr1b, first identified in the human brain [53], has been shown to function predominantly in the nucleus, akin to other FET protein family members [31, 36, 37]. Therefore, a key finding of the present study is that Ersr1b also functions in the cytoplasm to regulate mRNA translation (Figures 3–5). We previously demonstrated that *pou5f3* mRNA becomes translationally active as its RNA granules transition from a solid- to liquid-like state [23]. This transition coincided with the colocalization of Ewsr1b foci with *pou5f3* RNA granules (Figures 3E and 4A-B). Knockdown of Ewsr1b prevented this phase transition, inhibiting *pou5f3* mRNA translation, and reduced protein accumulation (Figures 4 and S4B), indicating that Ewsr1b is necessary for translational activation of dormant *pou5f3* mRNA. Additionally, the translation of second-wave mRNAs, such as *rbbp9* and *gmnn* (Figure 5D and E), was similarly diminished in Ewsr1b-deficient embryos (Figures 4E), suggesting that Ewsr1b broadly promotes the second mRNA translation wave. Consistently, Ewsr1b knockdown caused severe developmental defects beginning at an early stage (Figure 4G), underscoring the functional importance of Ewsr1b-mediated translational regulation in embryogenesis. Thus, Ewsr1b synthesized during the first-wave functions as a master regulator that triggers subsequent mRNA translation and early developmental processes.

FET proteins are generally soluble in the nucleus, but their mislocalization to the cytoplasm has been linked to pathological aggregate formation in neurodegenerative diseases [35, 54–57]. Our finding that Ewsr1b regulates maternal mRNA translation and early embryonic development (Figures 4 and S4) reveals previously unknown physiological roles for Ewsr1 and highlights their developmental relevance.

The question remains: how does Ewsr1b promote *pou5f3* mRNA translation? Ewsr1b is composed largely of IDRs (Figure 3A), which are unstructured domains, known to drive liquid–liquid phase separation, a process critical for the formation and regulation of membrane-less compartments, including RNA granules [34, 58–62]. Indeed, GFP-Ewsr1b exhibited liquid-like behavior in embryonic cells (Figure S11). Notably, zebrafish Ddx3xb has been shown to form liquid droplets through its IDR and enhance maternal mRNA translation around 4 hpf [63], and the eukaryotic translation initiation factor eIF4B has similarly been shown to self-assemble via IDRs and potentially promote mRNA translation [64]. Collectively, these observations support a model in which Ewsr1b directly binds to *pou5f3* RNA granules and induces a phase transition from a solid- to liquid-like state, thereby activating mRNA translation. Future studies should aim to elucidate the molecular organization of embryonic RNA granules and define how translational machinery assembles within them to initiate efficient mRNA translation.

### Alternative 3′UTRs of *ewsr1b* mRNA and multifunctionality of Ewsr1b

Our analyses revealed that Ewsr1b colocalizes with Pou5f3 protein in the nucleus (Figures 4C and S3). A previous study has shown that human EWSR1 regulates the transcriptional activity of Oct4/Pou5f1, a homolog of zebrafish Pou5f3, in the nucleus [65], suggesting that Ewsr1b may similarly influence Pou5f3-mediated transcription in the nucleus of zebrafish embryos. Supporting this, nuclear depletion of Ewsr1b reduced Pou5f3 accumulation and led to developmental defects at later stages (Figures 6D, E, and S7). These results indicate that Ewsr1b performs multiple functions during development, including translation regulation in the cytoplasm and may support protein stability and transcriptional activity in the nucleus.

Importantly, the temporal and spatial functions of Ewsr1b appear to be determined by the length of the 3′UTRs in its mRNA isoforms. The cytoplasmic form of Ewsr1b, which promotes mRNA translation, is synthesized during the first wave from *ewsr1b*-3′Short mRNA, whereas the nuclear-localized form is synthesized during the second translation wave from *ewsr1b*-3′Long mRNA (Figures 5–7). These distinct 3′UTRs are likely generated through alternative cleavage and polyadenylation, a widespread mechanism across cell types [66, 67]. Alternative 3′UTRs can modulate mRNA stability, localization, and translation efficiency by altering regulatory element composition [68–70]. In some cases, 3′UTRs can even serve as “nurturing niches” by recruiting proteins that cotranslationally regulate the localization and function of nascent proteins [69, 71, 72]. We found that Importin β1 binds to the long 3′UTR of *ewsr1b* mRNA, interacts with Ewsr1b in the cytoplasm and nucleus, and transports Ewsr1b in the nucleus (Figure 7A–E and S9). Based on these findings, we propose a model in which the long 3′UTR of *ewsr1b* mRNA recruits Importin β1, which cotranslationally interacts with newly synthesized Ewsr1b and facilities its nuclear import. In contrast, the absence of Importin β1 causes Ewsr1b translated from the short 3′UTR *ewsr1b* mRNA isoform to remain in the cytoplasmic.

One of the most unexpected findings of this study is that *ewsr1b* mRNA translated during the first-wave harbors a short 3′UTR comprising only 16 nucleotides (Figures 5–7). Other first-wave mRNAs also include variants with similarly short 3′UTRs (Figure S5A), suggesting that truncated 3′UTRs may represent a general strategy for the translational activation of dormant maternal mRNAs during the earliest stage of development. In general, short forms of alternative 3′UTRs increase mRNA translation efficiency due to deletion of cis-acting elements that repress translation [68, 73]. Our results uncovered a previously unrecognized role of alternative 3′UTRs that the short 3′UTR tightly represses translation before fertilization and rapidly initiates protein synthesis after fertilization, which induces sequential waves of mRNA translation crucial for whole embryonic development. HuR has been shown to regulate translation of target mRNAs [74]. We found that HuR binds *ewsr1b*-3′Short mRNA and that proteasome-mediated HuR degradation triggers translation of *ewsr1b*-3′Short mRNA immediately after fertilization (Figure 7F-I). Future studies should aim to identify the regulatory pathways that cause proteasome-mediated HuR degradation postfertilization. In summary, our findings provide fundamental insights into how complex biological processes are orchestrated at the posttranscriptional level and would help in understanding the cellular and molecular systems that drive diverse multilayered, long-term processes in a wide range of organisms.

### Limitations of the study

In this study, we discovered previously unknown principles for the control of mRNAs and embryonic development by focusing on Ewsr1b, Syncrip and HuR proteins. However, RNA-binding proteins have been shown to function cooperatively with many molecules in ribonucleoprotein (RNP) complexes [3, 34, 62] as observed in our mass spectrometry analysis. Comprehensive analyses for the RNP complexes will be needed in future investigations. Although we identified tens of mRNAs that increase translation rate in the earliest stages of development, the numbers of mRNAs in the distinct waves were not high possibly due to a limitation in the sensitivity of this analysis, and many mRNAs would increase translation rate in these periods. In addition, Ewsr1b, Syncrip and HuR target a wide range of mRNAs [75–77] and potentially regulate the translation of many mRNAs. These issues will be addressed in future studies.

## STAR METHODS

### KEY RESOURCES TABLE

A key resources table is provided in the latter part of this paper.

## RESOURCE AVAILABILITY

### Lead contact

Further information and requests for resources and reagents should be directed to and will be fulfilled by the lead contact, Tomoya Kotani.

### Materials availability

Plasmids and antibodies generated in this study are available from the lead contact upon completion of a Materials Transfer Agreement, but there are restrictions to the availability of antibodies due to the lack of an external centralized repository for its distribution and our need to maintain the stock.

### Dara availability

RNA sequencing data have been deposited in the Dryad under the dataset DOI: 10.5061/dryad.bzkh189q6. Information or requests for biological and chemical resources and reagents should be directed to the lead contact.

## EXPERIMENTAL MODEL AND SUBJECT DETAILS

### Animal husbandry

Adult zebrafish were maintained at 28°C on a 14-h light/10-h dark cycle under standard conditions. Embryos were obtained via natural spawning and staged according to hpf and morphological criteria. All embryos used in this study were collected prior to 24 hpf, during which sex is indeterminate and unlikely to influence the biological processes examined. All animal experiments in this study were approved by the Committee on Animal Experimentation, Hokkaido University.

## METHOD DETAILS

### Oocyte isolation and embryo collection

Zebrafish ovaries were dissected from adult females in Ringer’s solution (116 mM NaCl, 2.9 mM KCl, 1.8 mM CaCl_2_, and 5 mM HEPES; pH 7.2). Oocytes were manually isolated from ovaries using forceps under a dissecting microscope. Embryos were cultured in E3 medium (5 mM NaCl, 0.17 mM KCl, 0.33 mM CaCl_2_, and 0.33 mM MgSO_4_) and collected at defined developmental stages for subsequent experiments.

### Antibody production

A partial sequence encoding zebrafish Ewsr1b (residues 1–329) was amplified via PCR using gene-specific primers (*ewsr1b*-ORF-f1 and *ewsr1b*-ORF-r2) and cloned into the pET21 vector for expression of a histidine (His)-tagged recombinant protein. Protein expression was induced in *E. coli*, and the product was purified via SDS-PAGE followed by electroelution in Tris-glycine buffer without SDS. The protein was then dialyzed against 1 mM HEPES (pH 7.5), lyophilized, and used to immunize two mice. Antisera were affinity-purified using the recombinant His-tagged Ewsr1b protein electroblotted onto a membrane (Immobilon; EMD Millipore, IPVH00010). Primer sequences are listed in Table S3.

### Immunoblotting

Zebrafish embryos were obtained via natural spawning and staged according to hpf and morphological criteria. Embryos were homogenized in an equal volume of ice-cold extraction buffer [EB; 100 mM β-glycerophosphate, 20 mM HEPES, 15 mM MgCl_2_, 5 mM EGTA, 1 mM dithiothreitol, 100 μM (p-amidinophenyl) methanesulfonyl fluoride, and leupeptin (3 μg/mL); pH 7.5], supplemented with 0.04 U/µL recombinant RNase inhibitor (Takara, 2313A). Lysates were centrifuged at 15,000 rpm and 4°C for 10 min, and supernatants were collected as crude extracts. Proteins were separated by SDS-PAGE, transferred onto Immobilon membranes, and probed with primary antibodies; mouse anti-Syncrip antibody (1:1,000, hnRNP Q; Santa Cruz Biotechnology, I8E4; sc-56703), rabbit anti-Syncrip antibody (1:1,000, Proteintech, 14024-1-AP), rabbit anti-Rpl11 antibody (1:1,000, Abcam, ab79352), rabbit anti-Pou5f3 antibody [23] (1:100), mouse anti-GFP antibody (1:1,000, Roche, 11814460001), mouse anti-Ewsr1b antibody (1:100, present study), rabbit anti-Importin β1 antibody (1:1,000, Proteintech, 10077-1-AP), and mouse anti-HuR antibody (1:1000, Santa Cruz Biotechnology; sc-5261). Antibodies were diluted in blocking buffer (4% skim-milk and 0.1% Tween 20 in TBS buffer). Signal intensities were quantified using ImageJ software.

### RT-PCR

Total RNA from oocytes and embryos were extracted using TRIzol reagent (Invitrogen, 15596-018) following the manufacturer’s protocol. Reverse transcription was performed using the FastGene Scriptase Ⅱ cDNA Synthesis Kit (FastGene, NE-LS64) with oligo (dT) primers and 2.5 µg of total RNA. RT-PCR was performed using gene-specific primers targeting *syncrip* (*syncrip*-ORF-f1 and *syncrip*-ORF-r1), *ewsr1a* (*ewsr1a*-ORF-f1 and *ewsr1a*-ORF-r1), *ewsr1b* (*ewsr1b*-ORF-f1 and *ewsr1b*-ORF-r2), and *β-actin* (*β-actin*-ORF-f1 and *β-actin*-ORF-r1). Primer sequences are provided in Table S3.

### Whole-mount *in situ* hybridization

RNA probes for *in situ* hybridization were synthesized via PCR amplification of gene-specific regions for *pou5f3* (*pou5f3*-f1 and *pou5f3*-3’ UTR-r1), *ewsr1b* (ORF: *ewsr1b*-ORF-f1 and *ewsr1b*-ORF-r1; 3′UTR: *ewsr1b*-ORF-f2 and *ewsr1b*-3′UTR-r1), and *gfp* (*gfp*-f1 and *gfp*-r1). Amplified fragments were cloned into the pGEM-T Easy Vector (Promega, A1360), and plasmids were linearized using appropriate restriction enzymes. DIG- and fluorescein-labeled RNA probes were synthesized using SP6 or T7 RNA polymerase and DIG or fluorescein RNA Labeling Mix (Roche, 11277073910 or 11685619910).

Zebrafish embryos were fixed with 4% paraformaldehyde (PFA) in PBS overnight at 4°C. Chorions were removed in PBS using forceps, and embryos were treated with 100% methanol overnight at −20°C. After rehydration and washing in PBS containing 0.1% Tween 20, embryos were incubated for 5 min at 55°C in hybridization buffer without RNA (HYB-; 50% formamide, 5× SSC, and 0.1% Tween 20). Prehybridization was performed in HYB+ (HYB- supplemented with 5 mg/mL torula RNA and 50 µg/mL heparin) for 1 h at 55°C, followed by overnight hybridization at 65°C with RNA probes in HYB+. Embryos were then sequentially washed at 65°C in HYB- (30 min), 2× SSCT (2 x 30 min), and 0.2× SSCT (2 x 30 min).

Embryos were incubated in maleic acid buffer (150 mM maleic acid and 100 mM NaCl; pH 7.5) containing 0.1% Tween 20 (MBST) for 5 min, followed by blocking in 2% blocking reagent (Roche, 10447200) in MBST for 1 h. They were then incubated overnight at 4°C with anti-DIG-HRP antibody (1:700 dilution in blocking buffer), washed with MBST and PBS, and developed using tyramide-Fluorescein (Akoya Biosciences, NEL701A001KT) for 15 min at room temperature (RT) according to the manufacturer’s instructions. After washing and mounting in VECTASHIELD Mounting Medium with DAPI (Vector Laboratories, Inc, H1200), the samples were observed under a LSM980 confocal microscope (Zeiss).

To simultaneously detect the ORF and 3′UTR of *ewsr1b* mRNA following a previously reported procedure [23], embryos were hybridized using the fluorescein-labeled antisense RNA probe for *ewsr1b* 3′UTR and the DIG-labeled antisense RNA probe for *ewsr1b* ORF, followed first by incubation with anti-Fluorescein-HRP antibody (1:300 dilution) for 3 h and then by incubation with tyramide-Cy3 (Akoya Biosciences, NEL704A001KT), according to the manufacturer’s instructions. Subsequently, the embryos were incubated in 1% H_2_O_2_ in methanol for 30 min to inactivate HRP. After rehydration and washing with PBS, the embryos were incubated with anti-DIG-HRP antibody (1:700 dilution) for 3 h, followed by incubation with tyramide-Fluorescein (Akoya Biosciences, NEL701A001KT) following to the manufacturer’s instructions. The embryos were mounted and subjected to microscopy as described above. The distance between nuclear centers and *ewsr1b* mRNA signal localization was quantified using ImageJ.

### Immunofluorescence

Whole-mount immunofluorescence was performed as described previously [78]. Zebrafish embryos were fixed in 4% PFA in PBS overnight at 4°C. Following dechorionation in PBS, the embryos were treated with 100% methanol and stored at −20°C overnight. Rehydration was performed sequentially in 50% methanol/PBS and PBS alone, followed by two washes in PBTD buffer (0.1% Tween-20 and 1% DMSO in PBS) for 5 min per wash. The embryos were blocked in blocking buffer (1% bovine serum albumin, 10% new-born calf serum, and 0.3 M glycine in PBTD) for 1 h at RT, followed by incubation overnight at 4°C with primary antibodies diluted in blocking buffer. The following antibodies were used: rabbit anti-Syncrip (1:200), mouse anti-GFP (1:200), rabbit anti-Pou5f3 (1:200), mouse anti-Ewsr1b (1:50), and rabbit anti-Importin β1 (1:200). After the embryos were washed twice in PBTD (5 min per wash), they were incubated for 2 h at RT in fluorochrome-conjugated secondary antibodies (1:200 in blocking buffer). From this step onward, all procedures were conducted in the dark. After two additional PBTD washes (5 min per wash), the embryos were mounted in VECTASHIELD Mounting Medium and observed using an LSM980 confocal microscope. Signal intensity and particle counts were quantified using ImageJ.

For simultaneous detection of protein and mRNA, embryos were first hybridized with the DIG-labeled RNA probe as described above. After hybridization, protein detection was performed using primary and secondary antibodies according to the immunofluorescence protocol. After washing, the embryos were incubated with anti-DIG-HRP antibody, and RNA signals were detected using the tyramide signal amplification method, as described in the “Whole-mount *in situ* hybridization section”.

### IP/RT-PCR

Crude extracts from zebrafish embryos were incubated overnight at 4°C with mouse anti-Syncrip, mouse anti-GFP, rabbit anti-Rpl11, or rabbit anti-Importin β1 antibody in the presence of protein A Mag Sepharose beads (GE Healthcare). Mouse IgG or rabbit IgG of the same concentration was used as a control. After washing the beads, mRNA was extracted using TRIzol reagent (Invitrogen) and reverse transcribed using the FastGene Scriptase Ⅱ cDNA Synthesis Kit (FastGene) with oligo(dT) primers. RT-PCR was performed using gene-specific primer sets for *syncrip* (*syncrip*-ORF-f1 and *syncrip*-ORF-r2), *pou5f3* (*pou5f3*-3′UTR-f1 and *pou5f3*-3′UTR-r1), *α-tubulin* (*α-tubulin*-f1 and *α-tubulin*-r1), *β-actin* (*β-actin*-ORF-f2 and *β-actin*-ORF-r1), *ewsr1b* ORF (*ewsr1b*-ORF-f1 and *ewsr1b*-ORF-r3), *ewsr1b* 3′UTR (*ewsr1b*-ORF-f2 and *ewsr1b*-3′UTR-r1), *rbbp9* (*rbbp9*-f1 and *rbbp9*-r1), and *gmnn* (*gmnn*-f1 and *gmnn*-r1). Primer sequences are listed in Table S3.

### Immunoprecipitation followed by mass spectrometry

Following immunoprecipitation with anti-Syncrip antibody, samples were separated via SDS-PAGE. Gels were silver-stained using the Silver Stain MS Kit (Wako Pure Chemical Industries, Ltd.). Each lane was divided into five regions and subjected to mass spectrometric analysis using the Orbitrap Exploris 240 system (Thermo Fisher Scientific). Protein identification was performed using Sequest HT (Thermo Fisher Scientific) with Proteome Discoverer 3.0 software (Thermo Fisher Scientific), referencing the zebrafish (*Danio rerio*) section of the UniProtKB database (2024-1-24; 3314 sequences with 1,619,742 residues). Identifications were validated using the Percolator algorithm based on q-values. Proteins with at least one unique peptide match were considered. GO analysis was performed using the DAVID Knowledgebase v2023q4 (https://david.ncifcrf.gov/).

### qPCR

Total RNA was extracted from zebrafish oocytes and embryos using TRIzol reagent. Reverse transcription was performed using HiScript III All-in-one RT SuperMix Perfect for qPCR (Vazyme, R333-01). Subsequently, qPCR was conducted using gene-specific primer sets targeting *syncrip* (*syncrip*-ORF-f1 and *syncrip*-ORF-r3), *ewsr1a* (*ewsr1a*-ORF-f1 and *ewsr1a*-ORF-r2), *ewsr1b* (*ewsr1b*-ORF-f1 and *ewsr1b*-ORF-r4), *pou5f3* (*pou5f3*-qPCR-f1 and *pou5f3*-qPCR-r1), *α-tubulin* (*α-tubulin*-f2 and *α-tubulin*-r1), *β-actin* (*β-actin*-ORF-f1 and *β-actin*-qPCR-r1), *rbbp9* (*rbbp9*-f2 and *rbbp9*-r1), *gmnn* (*gmnn*-f2 and *gmnn*-r1), *kazald2* (*kazald2*-f1 and *kazald2*-r1), and *cldne* (*cldne*-f1 and *cldne*-r1) with Power SYBR green PCR Master Mix (Applied Biosystems, 4367659) on a real-time PCR system, following the manufacturer’s instructions. Primer sequences are presented in Table S3.

### PAT assay

RNA ligation–coupled RT-PCR was performed according to an established method [79]. Total RNA (2 µg), extracted from pools of 50 zebrafish embryos, was ligated with 0.4 μg of the P1 anchor primer in a 10-μL reaction using T4 RNA ligase (New England Biolabs) for 30 min at 37°C. Ligase was inactivated by heating at 92°C for 5 min. Subsequently, 5 μL of the ligation product was reverse transcribed (10-μl reaction) using the FastGene Scriptase Ⅱ cDNA Synthesis Kit with a P1′ primer. Next, 2 μL of cDNA was used for the first round of PCR (20 cycles, with a total volume of a 25 μL) with the P1′ primer and gene-specific primers targeting *pou5f3* (*pou5f3*-PAT-f1), *ewsr1b*-3′Long (*ewsr1b*-3′UTR-Long-PAT-f1), or *ewsr1b*-3′Short (*ewsr1b*-3′UTR-Short-PAT-f1). The first-round PCR product (1 µL) was then used for second-round PCR (35 cycles, with a total volume of 25 μL) with the P1′ primer and gene-specific primers targeting *pou5f3* (*pou5f3*-PAT-f2), *ewsr1b*-3′Long (*ewsr1b*-3′UTR-Long-PAT-f2), or *ewsr1b*-3′Short (*ewsr1b*-ORF-f2). PCR products were cloned into the pGEM-T Easy Vector (Promega) and sequenced. Primer sequences are listed in Table S3.

### MO injection

MO sequences (Gene Tools, LLC) are presented in Table S3 as well as Figure 5A and Figure S4A. The *ewsr1b*-ATG-MO (ATG-MO) targeted the start codon of *ewsr1b* mRNA, whereas the *ewsr1b*-5mm-MO (5mm-MO) contained 5-nt mismatches. The *ewsr1b*-5′UTR-MO (5′UTR-MO) targeted the 5′UTR of *ewsr1b* mRNA, and *ewsr1b*-3′end-MO (3′end-MO) targeted the 3′-end sequences of *ewsr1b* mRNA. MOs were diluted to 4 ng/nL in injection buffer (0.2 M KCl and 0.3% phenol red), and 1 nl of the solution was injected into fertilized eggs using a Femtojet microinjector (Eppendorf).

### Hexanediol treatment

To disrupt liquid-phase separated structures, zebrafish embryos were incubated in 10% (w/v) 1,6-hexanediol in E3 medium for 20 min at 28℃ (hexanediol was dissolved and diluted in E3 solution before use). Subsequently, embryos were fixed in 4% PFA in PBS overnight at 4°C and analyzed via whole-mount *in situ* hybridization.

### Immunoprecipitation followed by RNA sequencing

Embryos at selected developmental stages were dechorionated in E3 solution and incubated in 100 µg/mL cycloheximide in E3 for 5 min at RT. The embryos were then homogenized in an equal volume of extraction buffer and centrifuged at 15,000 rpm and 4°C for 10 min. The supernatant was collected for further processing. For total RNA samples, TRIzol reagent was added directly to the lysate for extraction. For Rpl11-IP samples, embryonic extracts were incubated overnight at 4°C with anti-Rpl11 antibody, or the same volume of rabbit IgG (control), followed by incubation with protein A Mag Sepharose beads overnight at 4°C. After washing, RNA was extracted from the beads using the TRIzol reagent.

RNA libraries were prepared using the SMART-Seq mRNA LP Kit (Takara) and Unique Dual Index Kits (Takara) according to the manufacturer’s protocols. RNA sequencing was performed on the NovaSeq X Plus platform (Illumina) via 150-bp paired-end reads. The run configuration generated a data package comprising 40 million reads, with the total output spanning one sequencing unit. Differential expression analysis was performed following the workflow provided by Macrogen Inc.

### 3′RACE

3′RACE was performed using 3′ Full RACE Core kit (Takara) according to the manufacturer’s protocols. After reverse transcription with the Oligo dT-3sites Adaptor Primer and 1 µg of RNA, 5 µL of cDNA was used for the first round PCR (20 cycles, with a total volume of a 25 µL) with the 3sites Adapter Primer and gene-specific primer targeting *ewsr1b* (*ewsr1b*-PAT-f1). The first round PCR product (1 µL) was then used for second-round PCR (35 cycles, with a total volume of 25 µL) with the 3sites Adapter Primer and gene-specific primer targeting *ewsr1b* (*ewsr1b*-PAT-f2). PCR products were cloned into the pGEM-T Easy Vector (Promega) and sequenced. Primer sequences are listed in Table S3.

### Injection of mRNA, live imaging, and FRAP

To construct the GFP-tagged reporter plasmids (GFP-Ewsr1b Long-, Short-, and 96-3′UTR), the respective 3′UTR sequences were amplified via PCR using specific primer sets (long: *ewsr1b*-ORF-f1 and *ewsr1b*-3′UTR-r1; short: *ewsr1b*-ORF-f1 and *ewsr1b*-ORF-r1; 96: *ewsr1b*-ORF-f1 and *ewsr1b*-3′UTR-r2). The mRNAs were synthesized using the mMESSAGE mMACHINE SP6 Transcription Kit (Thermo Fisher Scientific), and purified transcripts were dissolved in nuclease-free distilled water. One nanoliter of mRNA solution (200 ng/µL) was injected into fertilized zebrafish eggs.

For live imaging, embryos were mounted and observed using an LSM980 confocal microscope. FRAP analysis was also performed under this microscope. A circular region of interest (∼5 µm in diameter) within the embryonic cytoplasm was bleached using 100% 488-nm laser pulse for 1 s. Recovery images were captured at 1-s intervals for a total of 10 s using 1.0% laser power.

### Importazole and MG132 injection

Importin β inhibition was performed by injecting 1 nL of a solution containing 100 µM importazole (nuclear transport receptor importin β inhibitor; Abcam), 0.2 M KCl, and 0.3% phenol red into fertilized eggs. The injection solution was prepared by diluting 1 mM importazole in DMSO stock solution. Proteasomal inhibition was performed by injecting 1 nL of a solution containing 10 µM MG132 (proteasomal inhibitor; Wako Pure Chemical Industries, Ltd.), 0.2 M KCl, and 0.3% phenol red into fertilized eggs.

The injection solution was prepared by diluting 100 µM MG132 in DMSO stock solution, and an equivalent volume of DMSO was used for control injections. Embryos injected with importazole or MG132 were collected at appropriate developmental stages and processed for downstream analyses.

### Statistical analyses

All quantitative data are presented as means ± standard deviations (SDs). Comparisons between two groups were performed using Student’s *t*-test. For comparisons among multiple groups, the Tukey–Kramer test was applied, as noted in the figure legends. A p-value p <0.05 was considered statistically significant.

## Acknowledgements

This work was supported by Grant-in-Aid for Scientific Research KAKENHI grant numbers 21H02398, 23K18104, 24K01952 and 25H02435 (to T.K.) from Japan Society for the Promotion of Science (JSPS) and was in part supported by Grant-in-Aid for JSPS Fellows from JSPS grant number 23KJ0019 (to K.S.). This work was partly performed in the Cooperative Research Project Program of the Medical Institute of Bioregulation, Kyushu University.

## Autor contributions

Conceptualization, K.S, and T.K.; Investigation; K.S, T.S, F.L, and A.S.; Writing, K.S, and T.K.; Supervision and Funding acquisition, T.K.

## Competing interests

The authors declare no competing interests.

## Notes

### Competing Interest Statement

The authors have declared no competing interest.

### Summary of Updates

Section on Results has been updated to clarify functions of Ewsr1b, Syncrip and HuR. Figures 1, 4, 5, 7 have been revised.

