## Supplementary figures for "Ewsr1b, Syncrip, HuR and alternative 3′UTRs organize sequential waves of translation to drive embryonic development"

**Figure S1. Syncrip expression decreases after fertilization at the mRNA and protein levels.** (A) Left: RT-PCR of *syncrip* and *β-actin* mRNAs in oocytes and embryos at 0, 3, and 6 hpf. Right: RT-qPCR of *syncrip* mRNA in embryos at 0, 3, and 6 hpf (means ± SD; n = 3). $*p < 0.05$; $**p < 0.01$; $***p < 0.001$ (Tukey–Kramer test). (B) Left: Immunofluorescence of Syncrip in embryos at 0, 3, and 6 hpf. DNA is shown in blue. Scale bars: 20 μm. Right: Quantification of Syncrip signals per 400 $\mu m^2$ in embryos at 0, 3, and 6 hpf (means ± SD; n = 12). Similar results were obtained from three independent experiments. NS, not significant; $**p < 0.01$; $***p < 0.001$ (Tukey–Kramer test). (C) STRING-based (https://string-db.org) interaction network of Syncrip-associated proteins. Edges represent predicted functional associations.

**Figure S2. Expression patterns of *ewsr1a* and *ewsr1b* mRNAs.** (A) Schematic domain structures of zebrafish Ewsr1a and Ewsr1b. SYGQ-rich, serine-tyrosine-glycine-glutamine–rich domain; RGG, arginine-glycine-glycine–rich domain; RRM, RNA recognition domain; ZnF, zinc finger domain; NLS, nuclear localization signal. (B) Left: RT-PCR of *ewsr1a*, *ewsr1b*, and *β-actin* mRNAs in oocytes and embryos at 6 hpf. Right: RT-qPCR of *ewsr1a* and *ewsr1b* mRNAs in oocytes (left) and embryos at 6 hpf (right) (means ± SD; n = 3). $******p < 0.000001$; $*******p < 0.0000001$ (Student's *t*-test). (C) Left: RT-PCR of *ewsr1b* mRNA in oocytes and embryos at 0, 3, and 6 hpf. Right: RT-qPCR of *ewsr1b* mRNA in embryos at 0, 3, and 6 hpf (means ± SD; n = 3). NS, not significant (Tukey–Kramer test).

**Figure S3. Ewsr1b interacts with Pou5f3 in early embryos.** (A) PLA detecting Ewsr1b and Pou5f3 (Ewsr1b-Pou5f3 PLA) in embryos at 0, 1, 2, and 3 hpf. DNA is shown in blue. Scale bars: 10 μm. (B) Quantification of PLA signals per 400 $\mu m^2$ in embryos at 0, 1, 2, and 3 hpf (means ± SD; n = 6). Similar results were obtained from two independent experiments. NS, not significant; $**p < 0.01$; $*******p < 0.0000001$ (Tukey–Kramer test).

**Figure S4. Ewsr1b knockdown inhibits *pou5f3* RNA granule phase change and causes developmental defects.** (A) Target sequences in the 5′UTR and ORF of *ewsr1b* mRNA for MOs: *ewsr1b*-ATG-MO (ATG-MO) and *ewsr1b*-5′UTR-MO (5′UTR-MO).

Notably, *ewsr1b*-5mm-MO (5mm-MO) contains 5-nucleotide mismatches (blue). (B) Left: Whole-mount *in situ* hybridization of *ewsr1b* mRNA in untreated (Cont), hexanediol-treated (Cont + HD), and ATG-MO-injected hexanediol-treated (ATG-MO + HD) embryos at 3 hpf. Bottom panels: Enlarged views of boxed areas. Scale bars: 20 µm (top) and 10 µm (bottom). Right: Quantification of *pou5f3* RNA granules per 100 $\mu m^2$ in embryos at 3 hpf (means ± SD; n = 12). Similar results were obtained from two independent experiments. NS, not significant; ****p < 0.0001; *****p < 0.00001 (Tukey–Kramer test). (C) RT-qPCR of *β-actin*, *kazald2*, and *cldne* mRNAs in uninjected embryos (WT) and those injected with ATG-MO (ATG) at 4 hpf (means ± SD; n = 3). Similar results were obtained from three independent experiments. *p < 0.05 (Student's *t*-test). (D) Percentage of embryos with developmental defects at 24 hpf (means ± SD; n = 6). ********p < 0.00000001 (Student's *t*-test). (E) Lateral views of wild-type embryos (WT) and those injected with 5′UTR-MO (5′UTR-MO) or 5′UTR-MO mixed with Short- and Long-3′UTR mRNAs (5′UTR-MO + mRNA; refer to Fig. 6) at 3, 6, 8 and 22 hpf. Bottom panels: GFP fluorescence images. Scale bars: 300 µm. Similar results were obtained from three independent experiments. (F) Percentages of normal and abnormal embryos at 22 hpf with (+) or without (-) 5′UTR-MO and mRNA injection.

**Figure S5. Syncrip and Ewsr1b regulate translation of target mRNAs.** (A) Tracks of 3′-end RNA sequencing plotted on the last exons of *ewsr1b*, *rrs1*, and *rps20* in embryos at 0 hpf. Arrows indicate short 3′UTR reads. (B) 3′RACE of *ewsr1b* mRNA in embryos at 0 hpf after IP using control IgG (IgG) or anti-Syncrip (α-Syncrip) antibody. Bottom: Sequence analysis of the 3′RACE products. (C) Left: Immunoblotting of GFP-Syncrip, GFP, Ewsr1b, and Rpl11 in uninjected embryos (−) and those injected with GFP (GFP) or GFP-Syncrip (GFP-Syncrip) at 6 hpf. Right: Quantification of immunoblotting (means ± SD; n = 3). Similar results were obtained from three independent experiments. NS, not significant; *p < 0.05; **p < 0.01 (Tukey–Kramer test). (D) Changes in read counts of *ewsr1b* and *pou5f3* from 0 to 4 hpf.

**Figure S6. Proximal 3′UTR region of *ewsr1b* is sufficient for nuclear localization of Ewsr1b.** (A) Top: Schematic diagrams of GFP-Ewsr1b Short- (Short-3′UTR mRNA), 96nt- (96nt-3′UTR mRNA), and Long-3′UTR (Long-3′UTR mRNA) mRNAs. Bottom:

Immunoblotting of uninjected embryos (−) and those injected with 96nt-3′UTR mRNA at 6 hpf after IP with anti-GFP antibody. (B) Localization of GFP-Ewsr1b translated from the three mRNA variants in embryos at 6 hpf. Similar results were obtained from two independent experiments. Scale bars: 20 μm.

**Figure S7. Ewsr1b synthesized from *ewsr1b*-3′Long mRNA is required for later-stage development.** (A) Lateral views of WT embryos and those injected with *ewsr1b*-3′end-MO (3′end-MO) at 22 and 46 hpf. Scale bars: 300 μm. Similar results were obtained from three independent experiments. (B) Percentages of morphological abnormalities at 22 hpf in WT and 3′end-MO-injected embryos.

**Figure S8. Subcellular localization of *ewsr1b* mRNA correlates with Ewsr1b localization.** (A) Schematic of dual *in situ* hybridization distinguishing *ewsr1b*-3′Long and *ewsr1b*-3′Short mRNAs. (B) Double *in situ* hybridization of ORF (green) and 3′UTR (red) for *ewsr1b* mRNA in embryos at 0 and 3 hpf. DNA is shown in blue. Left: High-resolution confocal images; right: enlarged views of boxed areas. Solid-line circles: *ewsr1b*-3′Long mRNAs; dotted-line circles: *ewsr1b*-3′Short mRNAs. (C) Left: Representative measurements from the nucleus center to *ewsr1b*-3′Long (white lines) and *ewsr1b*-3′Short mRNA (light blue dotted lines) at 3 hpf embryo. DNA is shown in blue. Right: Violin plot of distances from the nucleus center to signals of *ewsr1b*-3′Long (Long) and *ewsr1b*-3′Short (Short) mRNAs (means ± SD; n = 60). Similar results were obtained from three independent experiments. **p < 0.01 (Student's *t*-test). Scale bars: 20 μm.

**Figure S9. Importin β1 transports Ewsr1b in the nucleus.** (A) PLA of Ewsr1b and Importin β1 in embryos at 3 and 6 hpf. Top panel: high-resolution confocal images in embryos at 3 and 6 hpf. Right: Image of embryos at 3 hpf without anti-Ewsr1b antibody (No Ewsr1b 1st Ab). Bottom panel: Enlarged views of boxed areas in upper images. Similar results were obtained from two independent experiments. Scale bars: 20 μm. (B) Immunofluorescence of Importin β1 in uninjected embryos (Control) or embryos injected with Importazole at 3 hpf. DNA is shown in blue. Enlarged views of the boxed area with or without DNA staining are shown on the right side. Scale bars, 10 μm. (C) Quantification of average signal intensity in the nucleus per 25 $\mu m^2$. (means ± SD; n =

10). ***********$p < 0.00000000001$ (Student's *t*-test). (D) PLA of Ewsr1b and Importin β1 in uninjected embryos (Control) or embryos injected with Imporazole at 6 hpf. Similar results were obtained from two independent experiments. Scale bars, 10 µm. (E) Localization of GFP-Ewsr1b synthesized from GFP-Ewsr1b mRNA carrying Long-3′UTR in uninjected embryos (Control) or embryos injected with Importazole at 4 hpf. Similar results were obtained from two independent experiments. Scale bars, 10 µm.

**Figure S10. Proteasome activity is required for the first translation wave of *ewsr1b* mRNA.** (A) Immunofluorescence of Ewsr1b in uninjected embryos (Control) and those injected with DMSO or MG132 at 1 hpf. Similar results were obtained from two independent experiments. Scale bars: 20 µm. (B) Lateral views of WT embryos and those injected with MG132 at 2 and 5 hpf. Scale bars: 300 µm. Similar results were obtained from two independent experiments. (C) PAT assay for *ewsr1b*-3′Short mRNA in uninjected embryos (−) and those injected with DMSO or MG132 at 0 and 1 hpf. Similar results were obtained from two independent experiments. (D) Top: Immunoblotting of Rpl11 in uninjected embryos (-) and embryos injected with MG132 at 1 hpf after IP with anti-Rpl11 (α-Rpl11) antibody. Bottom: RT-PCR for *ewsr1b* mRNA ORF (*ewsr1b* ORF) and *ewsr1b* mRNA 3′UTR (*ewsr1b* 3′UTR). An asterisk indicates nonspecific band. Similar results were obtained from two independent experiments. (E) 3′RACE of *ewsr1b* mRNA in embryos at 0 hpf after IP using control IgG (IgG) or anti-HuR (α-HuR) antibody. Right: Sequence analysis of the 3′RACE products.

**Figure S11. Ewsr1b synthesized from *ewsr1b*-3′Short mRNA exhibits liquid-like properties.** (A) Left: Subcellular distribution of GFP-Ewsr1b synthesized from 3′UTR-Short mRNA in embryos at 6 hpf. Right: Time-lapse images of droplet fusion (arrows) in the boxed area. Similar results were obtained from three independent experiments. Scale bars: 10 µm. (B) Representative FRAP images of GFP-Ewsr1b before and 3-s after photobleaching. Similar results were obtained from three independent experiments. (C) Fluorescence recovery curves of GFP-Ewsr1b (means ± SD; n = 3). Similar results were obtained from three independent experiments.

### Figure S1

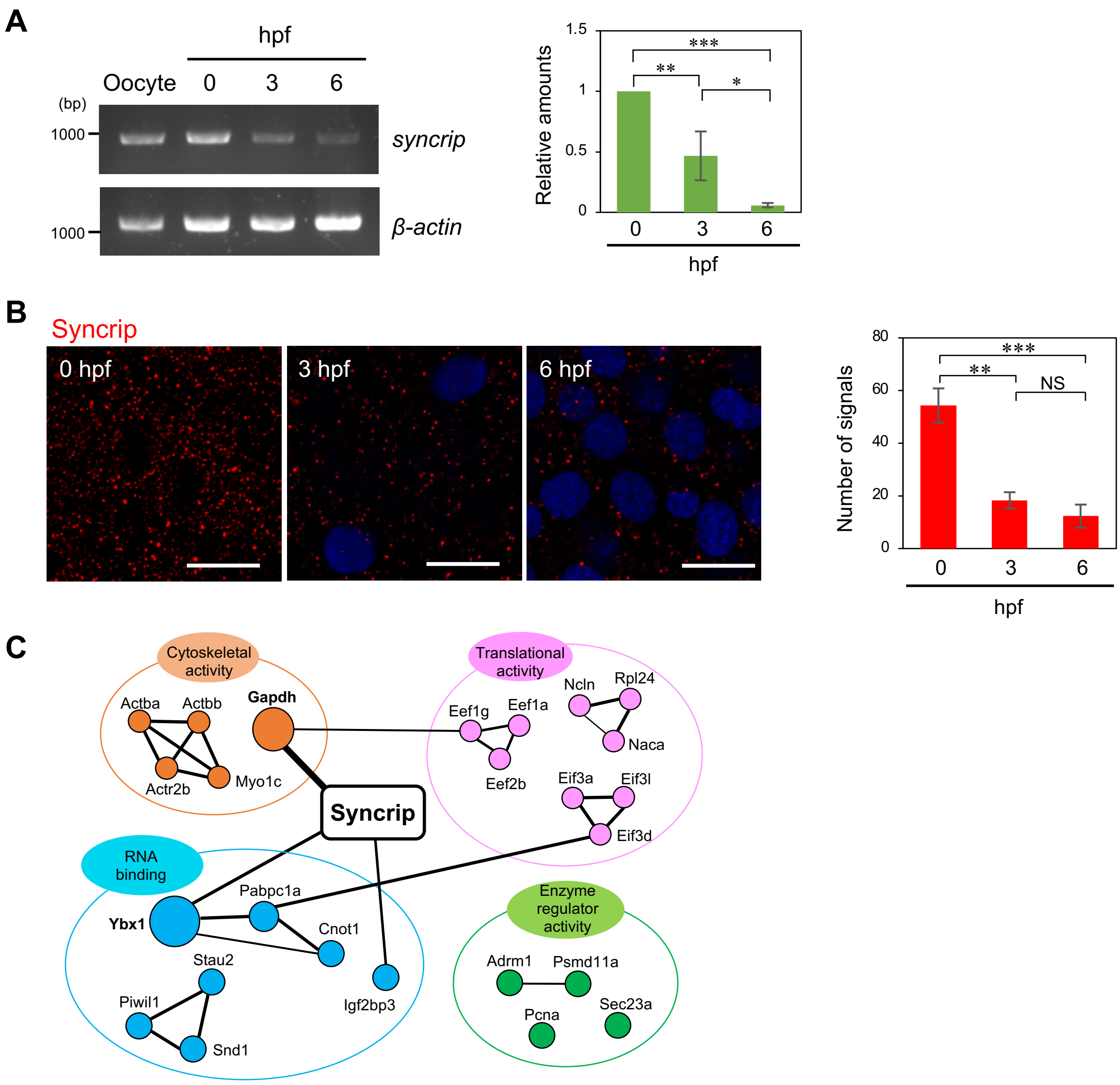
A
hpf
Oocyte 0 3 6
(bp)
1000
syncrip
1000
β-actin
Relative amounts
1.5
1
0.5
0
***
**
*
0 3 6
hpf
B
Syncrip
0 hpf
3 hpf
6 hpf
Number of signals
80
60
40
20
0
***
**
NS
0 3 6
hpf
C
Cytoskeletal activity
Actba
Actbb
Gapdh
Actr2b
Myo1c
Translational activity
Ncln
Rpl24
Eef1g
Eef1a
Naca
Eef2b
Eif3a
Eif3l
Eif3d
Syncrip
RNA binding
Ybx1
Pabpc1a
Cnot1
Stau2
Piwil1
Snd1
Igf2bp3
Enzyme regulator activity
Adrm1
Psmd11a
Pcna
Sec23a

Figure S2

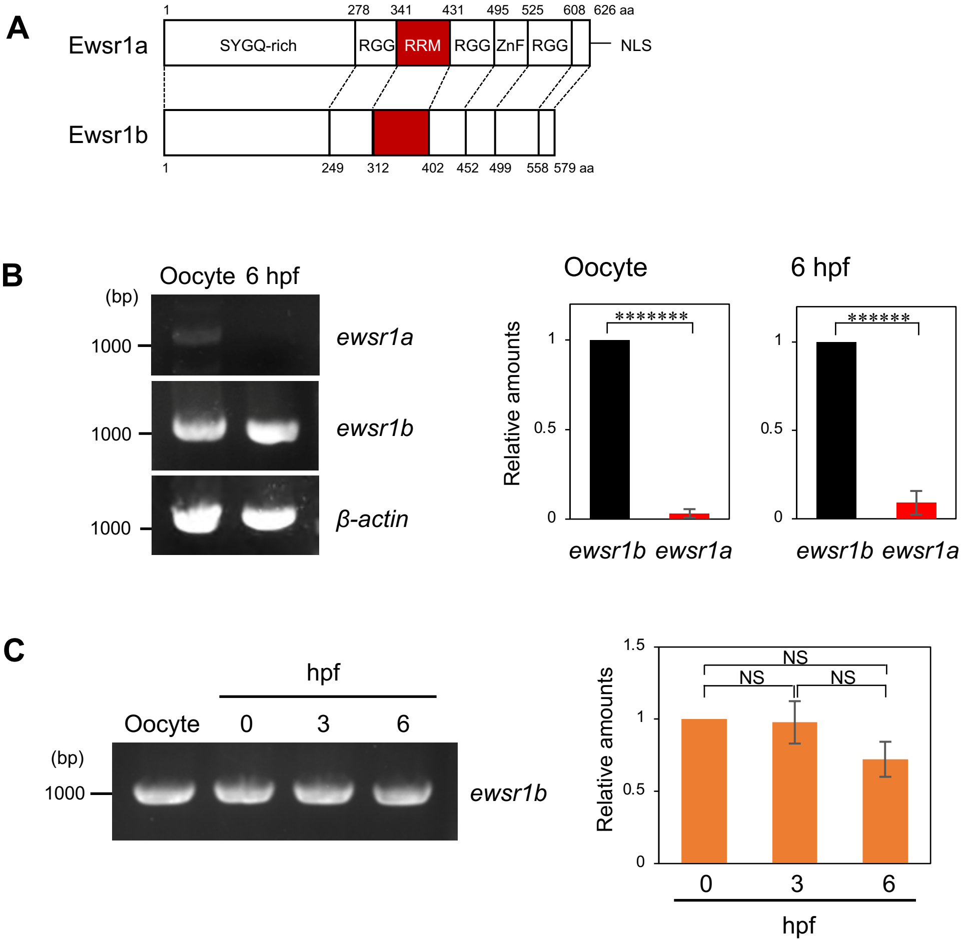
A
Ewsr1a
1
278
341
431
495
525
608
626 aa
SYGQ-rich
RGG
RRM
RGG
ZnF
RGG
NLS
Ewsr1b
1
249
312
402
452
499
558
579 aa
B
Oocyte
6 hpf
(bp)
1000
ewsr1a
ewsr1b
β-actin
Oocyte
6 hpf
Relative amounts
*******
******
1
0.5
0
ewsr1b
ewsr1a
C
hpf
Oocyte
0
3
6
(bp)
1000
ewsr1b
1.5
1
0.5
0
NS
NS
NS
Relative amounts
0
3
6
hpf

Figure S3

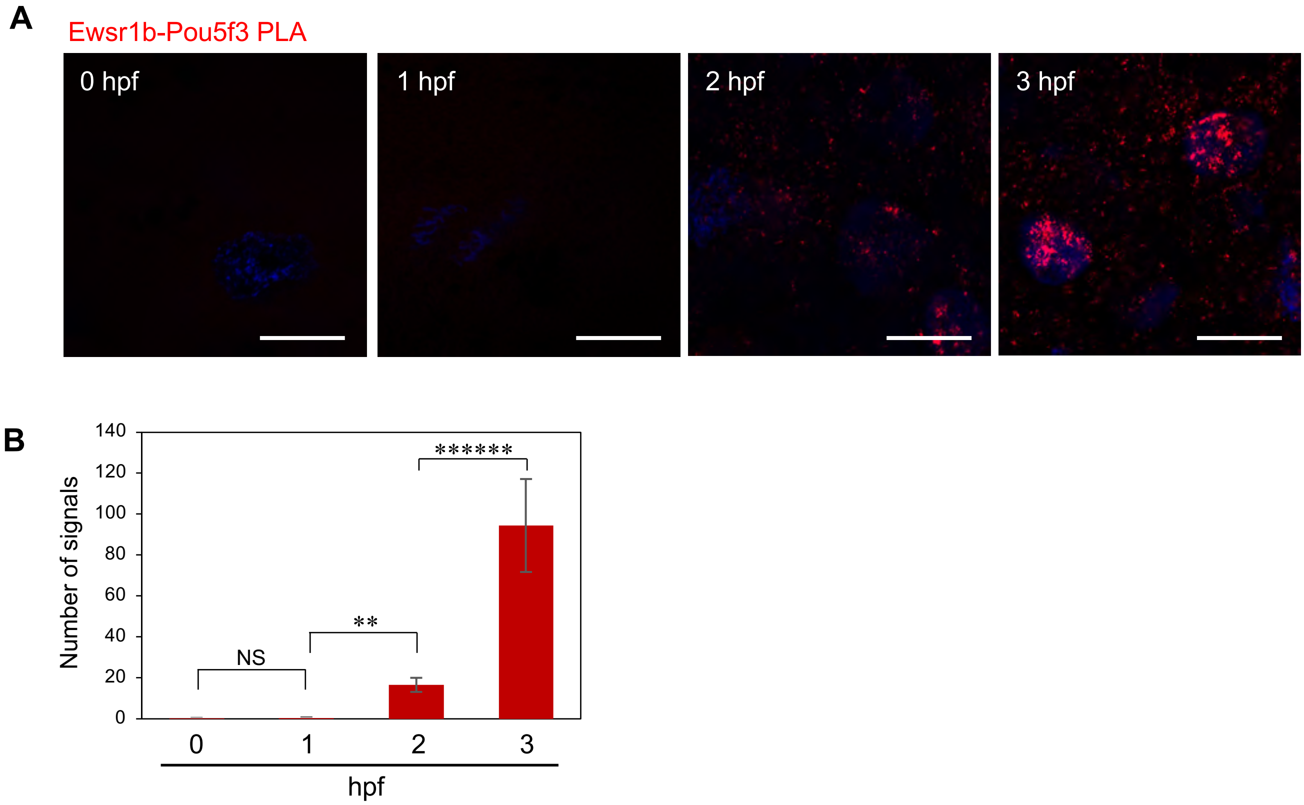
A
Ewsr1b-Pou5f3 PLA
0 hpf
1 hpf
2 hpf
3 hpf
B
Number of signals
140
120
100
80
60
40
20
0
NS
**
******
0
1
2
3
hpf

Figure S4

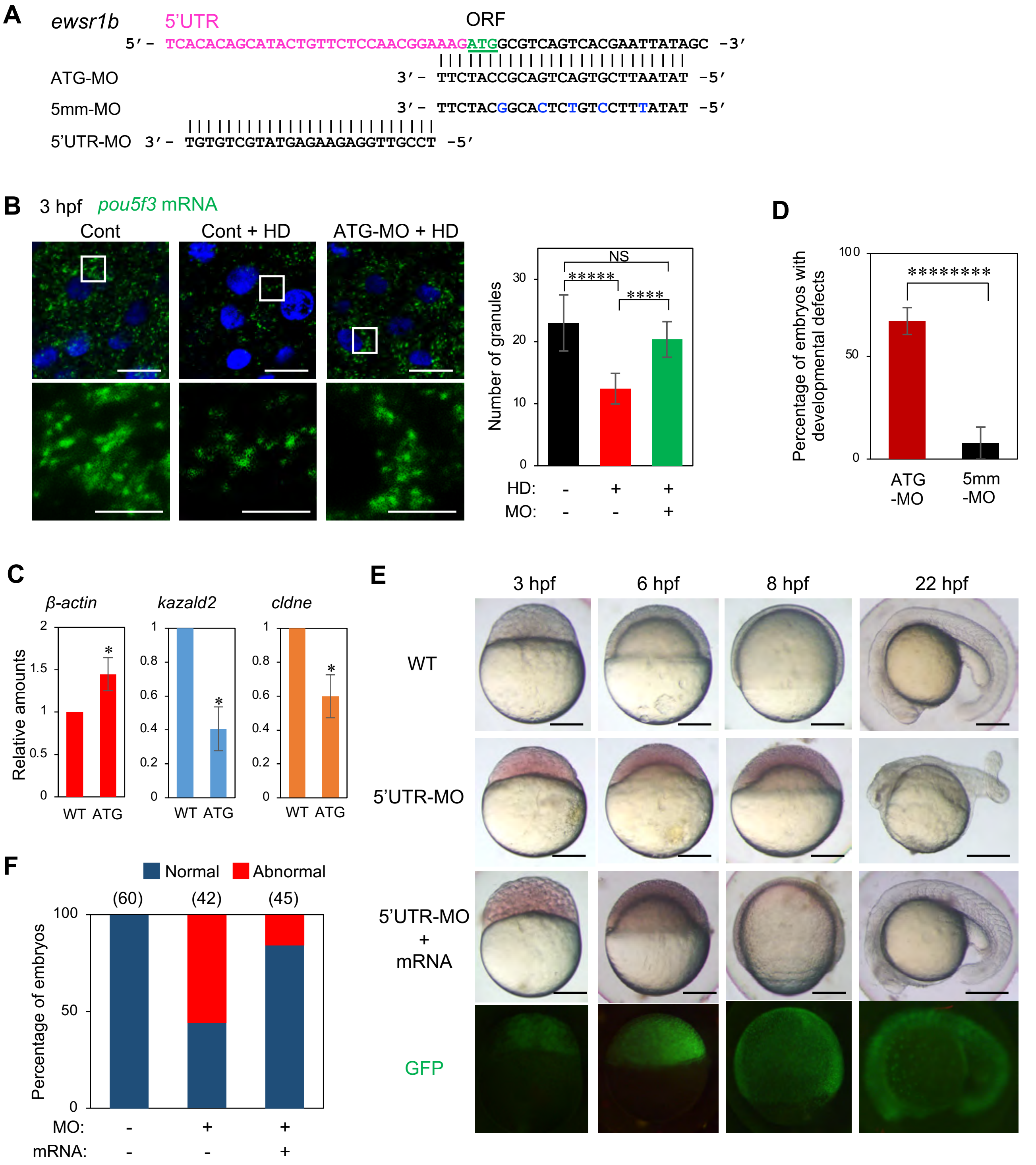

A
ewsr1b
5'UTR
ORF
5'- TCACACAGCATACTGTTCTCCAACGGAAAGATGGCGTCAGTCACGAATTATAGC -3'
ATG-MO
3'- TTCTACCGCAGTCAGTGCTTAATAT -5'
5mm-MO
3'- TTCTACGGCACTCTGTCCTTTATAT -5'
5'UTR-MO
3'- TGTGTCGTATGAGAAGAGGTTGCCT -5'
B
3 hpf
pou5f3 mRNA
Cont
Cont + HD
ATG-MO + HD
Number of granules
NS
*****
****
HD:
MO:
D
Percentage of embryos with developmental defects
********
ATG -MO
5mm -MO
C
β-actin
kazald2
cldne
Relative amounts
WT
ATG
E
3 hpf
6 hpf
8 hpf
22 hpf
WT
5'UTR-MO
5'UTR-MO + mRNA
GFP
F
Normal
Abnormal
(60)
(42)
(45)
Percentage of embryos
MO:
mRNA:

Figure S5

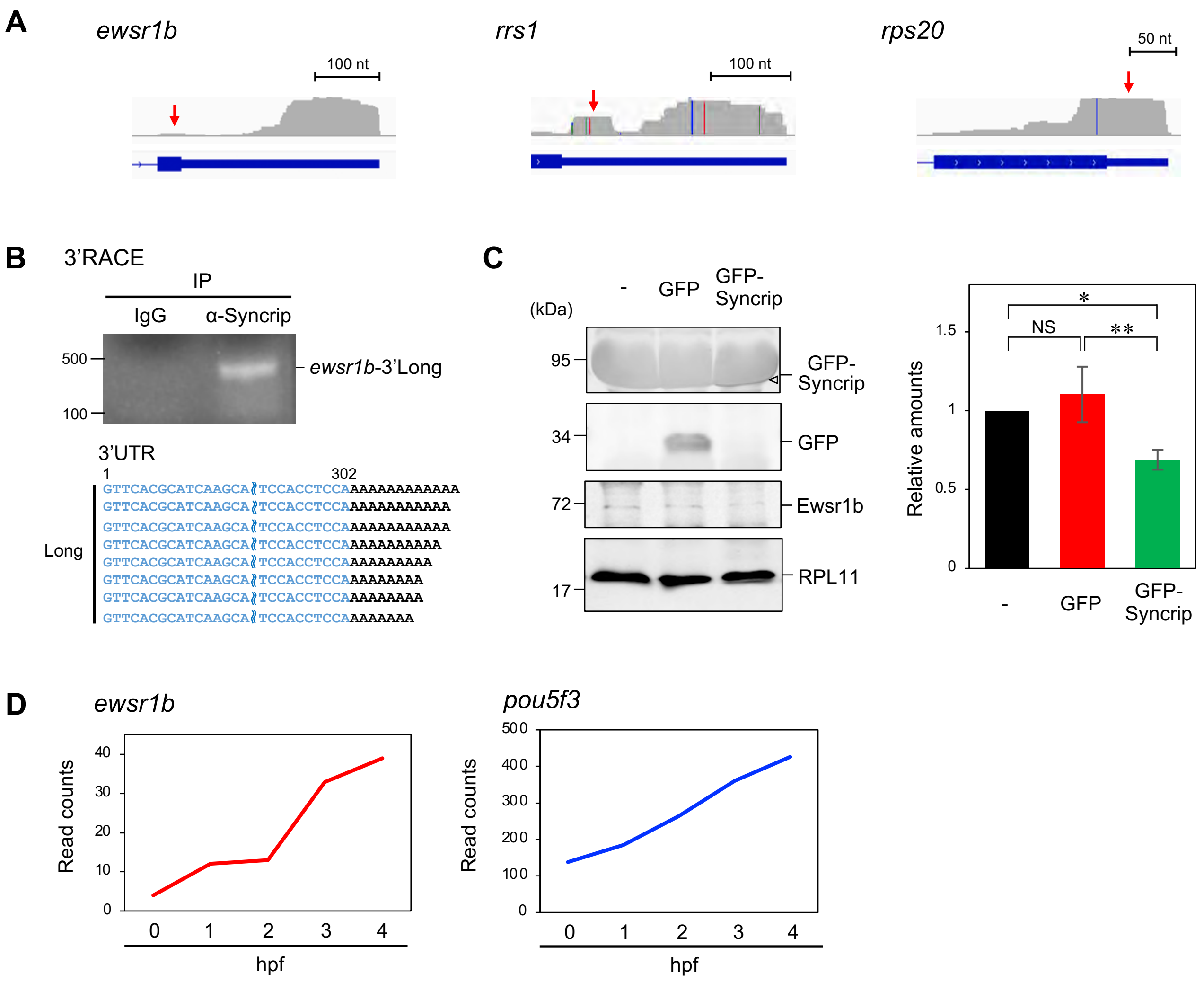
A
ewsr1b
100 nt
rrs1
100 nt
rps20
50 nt
B
3'RACE
IP
IgG
α-Syncrip
500
100
ewsr1b-3'Long
3'UTR
1
302
Long
GTTCACGCATCAAGCA TCCACCTCCAAAAAAAAAAAAA
C
(kDa)
-
GFP
GFP-Syncrip
95
34
72
17
GFP-Syncrip
GFP
Ewsr1b
RPL11
Relative amounts
1.5
1
0.5
0
NS
*
**
D
ewsr1b
pou5f3
Read counts
hpf
40
30
20
10
0
500
400
300
200
100
0 1 2 3 4

Figure S6

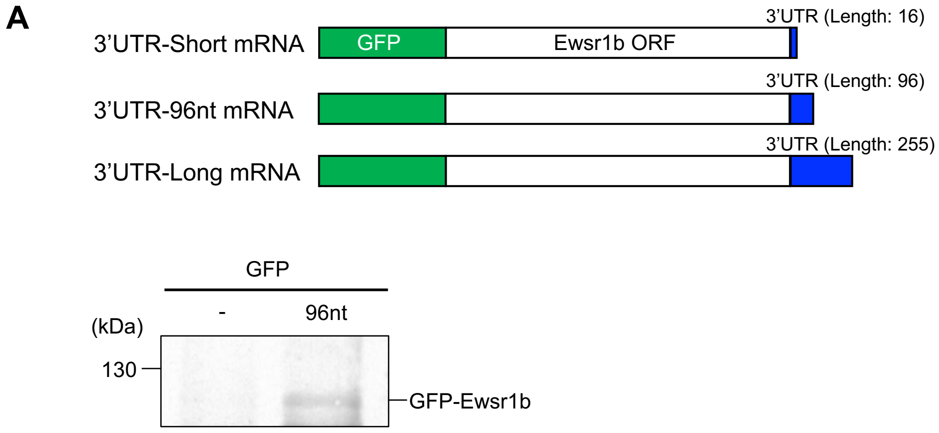
A
3'UTR-Short mRNA
GFP
Ewsr1b ORF
3'UTR (Length: 16)
3'UTR-96nt mRNA
3'UTR (Length: 96)
3'UTR-Long mRNA
3'UTR (Length: 255)
GFP
-
96nt
(kDa)
130
GFP-Ewsr1b

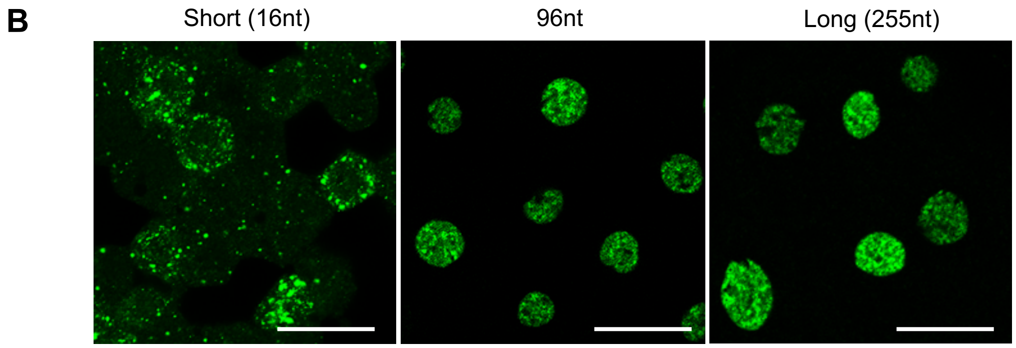
B
Short (16nt)
96nt
Long (255nt)

Figure S7

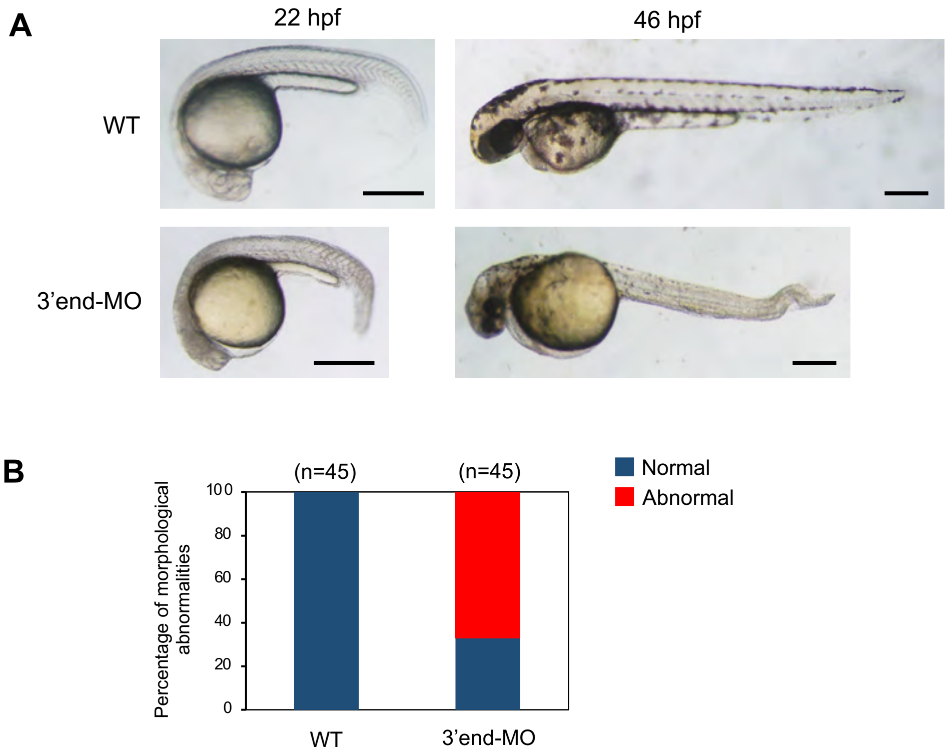

A
22 hpf
46 hpf
WT
3’end-MO
B
(n=45)
(n=45)
Normal
Abnormal
Percentage of morphological abnormalities
100
80
60
40
20
0
WT
3’end-MO

Figure S8

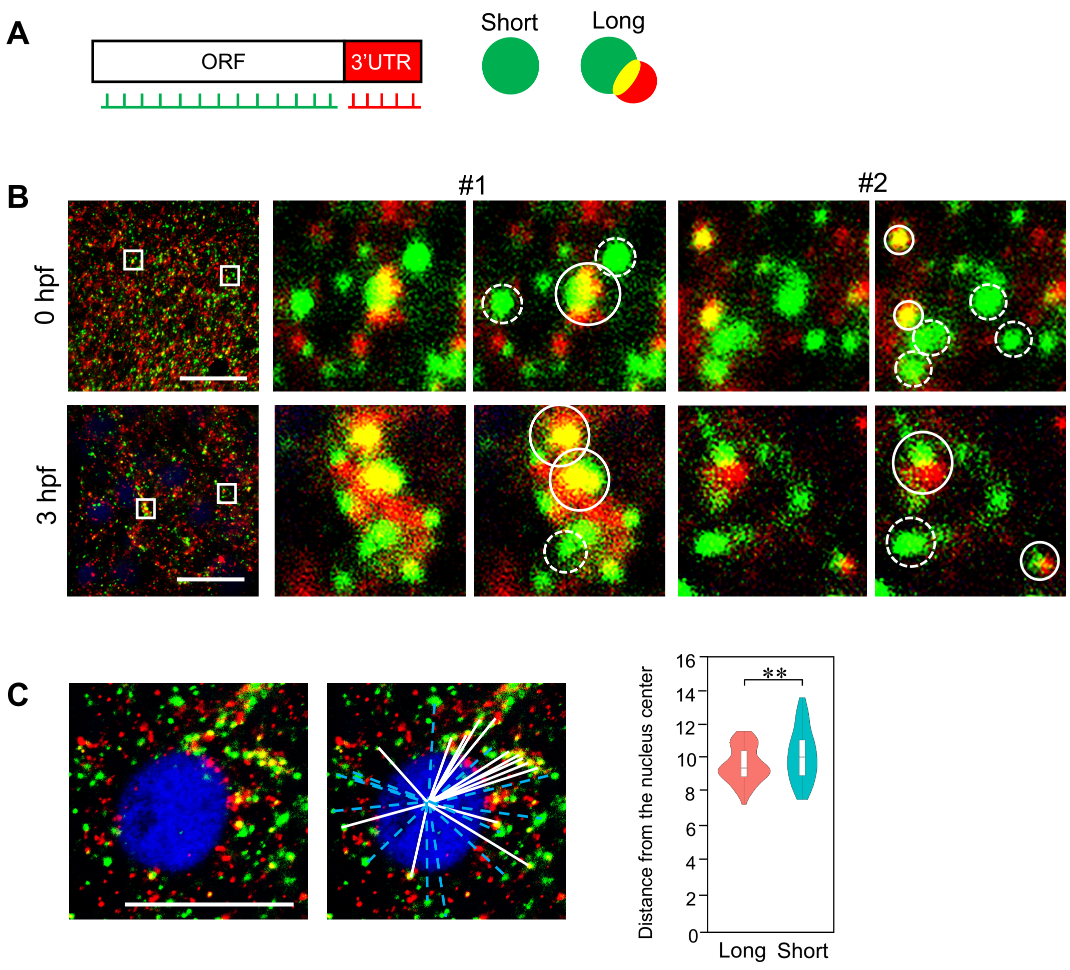
A
ORF
3'UTR
Short
Long
B
#1
#2
0 hpf
3 hpf
C
16
14
12
10
8
6
4
2
0
**
Distance from the nucleus center
Long
Short

Figure S9

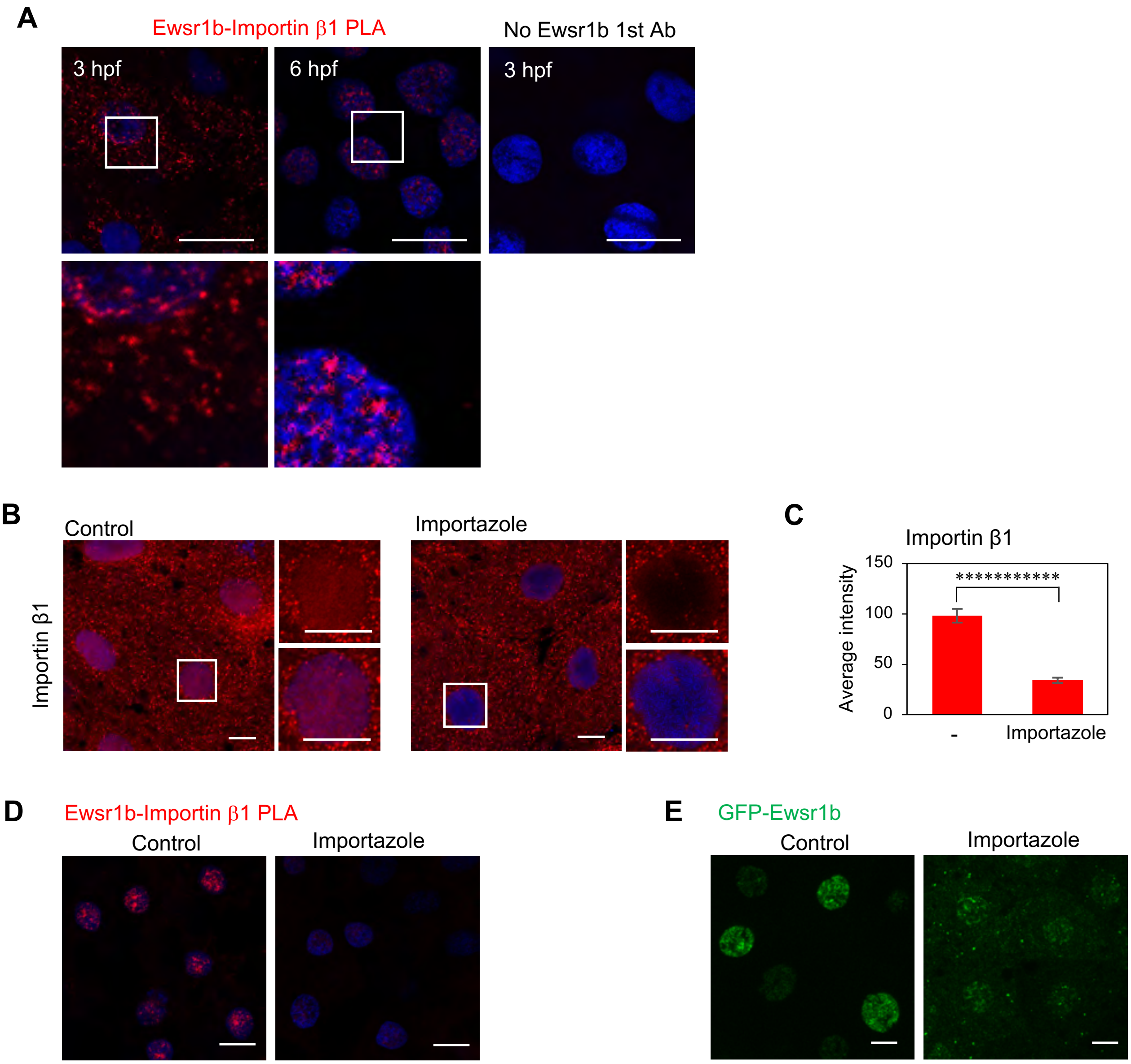
A
Ewsr1b-Importin β1 PLA
No Ewsr1b 1st Ab
3 hpf
6 hpf
3 hpf
B
Control
Importazole
Importin β1
C
Importin β1
Average intensity
150
100
50
0
***********
-
Importazole
D
Ewsr1b-Importin β1 PLA
Control
Importazole
E
GFP-Ewsr1b
Control
Importazole

Figure S10

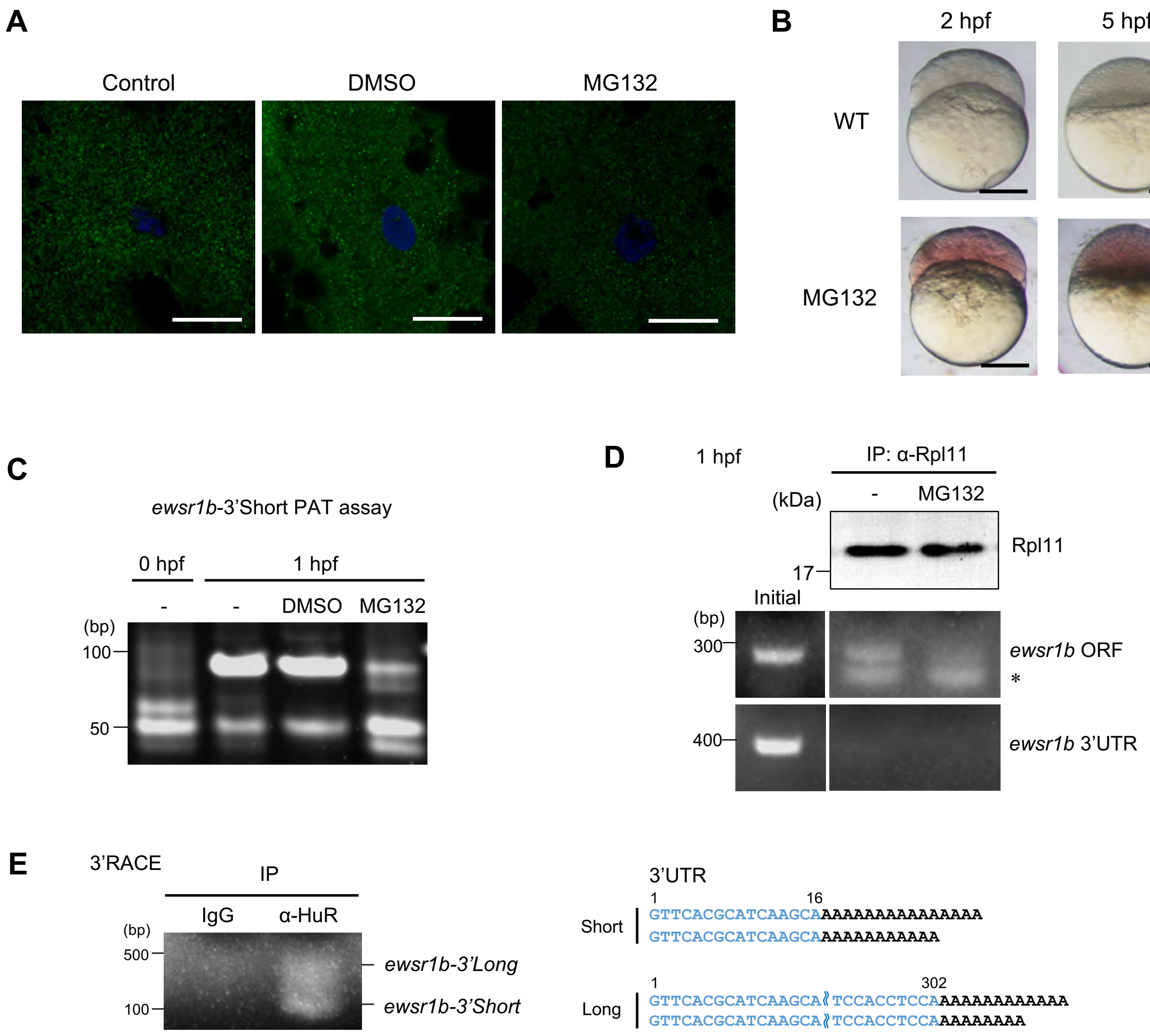

A
Control
DMSO
MG132
B
2 hpf
5 hpf
WT
MG132
C
ewsr1b-3'Short PAT assay
0 hpf
1 hpf
-
-
DMSO
MG132
(bp)
100
50
D
1 hpf
IP: α-Rpl11
(kDa)
-
MG132
Rpl11
17
Initial
(bp)
300
ewsr1b ORF
*
400
ewsr1b 3'UTR
E
3'RACE
IP
IgG
α-HuR
(bp)
500
100
ewsr1b-3'Long
ewsr1b-3'Short
3'UTR
1
16
Short
GTTCACGCATCAAGCAAAAAAAAAAAAAAAA
GTTCACGCATCAAGCAAAAAAAAAAAA
1
302
Long
GTTCACGCATCAAGCA≀TCCACCTCCAAAAAAAAAAAAA
GTTCACGCATCAAGCA≀TCCACCTCCAAAAAAAAA

Figure S11

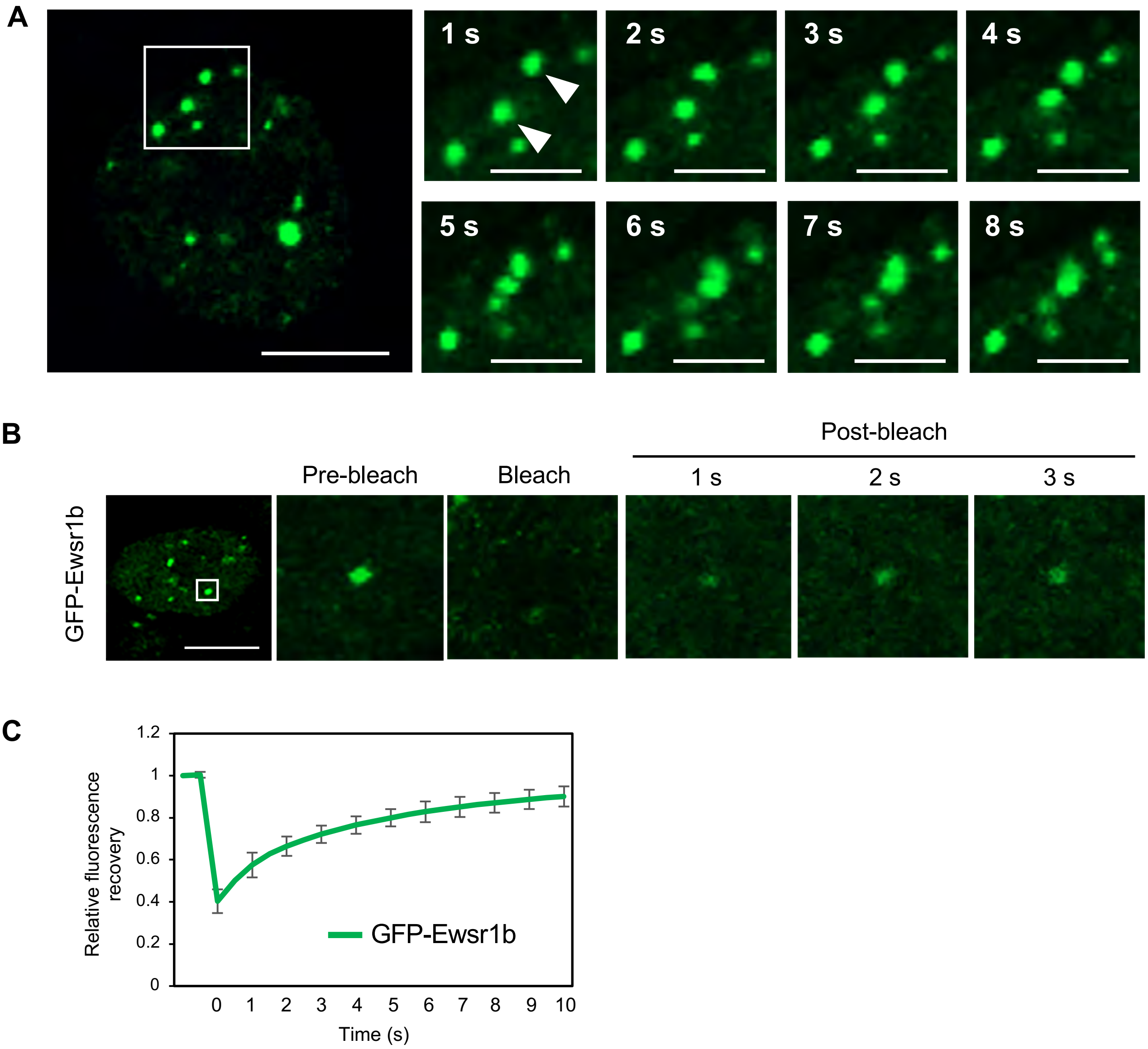
A
1 s
2 s
3 s
4 s
5 s
6 s
7 s
8 s
B
Post-bleach
Pre-bleach
Bleach
1 s
2 s
3 s
GFP-Ewsr1b
C
Relative fluorescence recovery
1.2
1
0.8
0.6
0.4
0.2
0
GFP-Ewsr1b
0 1 2 3 4 5 6 7 8 9 10
Time (s)
